# Versatile roles of Annexin A4 in ccRCC: impact on membrane repair, transcriptional signatures, and composition of the tumor microenvironment

**DOI:** 10.1101/2024.05.31.596161

**Authors:** Maximilian Wess, Manuel Rogg, Constance Gueib-Picard, Annika Merz, Anna L. Koessinger, Tobias Feilen, Grigor Andreev, Martin Werner, Ian J. Frew, Markus Grabbert, Oliver Schilling, Christoph Schell

## Abstract

Clear cell renal cell carcinoma (ccRCC) is the most prevalent type of renal malignant disease and is characterized by dismal prognosis in the metastasized setting. Invasive growth of cancer cells relates to high levels of compressive forces translating to relevant damage of the plasma membrane. However, functional implications of protein machineries required for plasma membrane repair in ccRCC are not yet completely elucidated. Given the membrane-associated localization of the large family of annexin proteins, we aimed for a global annotation of annexin proteins, which led to the identification of ANXA4 selectively expressed in cancer cells of ccRCC. Interestingly, ANXA4 showed context-dependent distinct localization patterns including the plasma membrane as well as the nuclear compartment/nuclear membrane. We investigated the functional role of ANXA4 in ccRCC employing genetic titration studies (knockdown, CRISPR/Cas9 knockout and overexpression) and identified impaired acute plasma membrane repair as well as invasive capability in conditions of reduced ANXA4 expression. Utilizing computational segmentation of the tumor microenvironment (TME) of ccRCC samples revealed that ANXA4 low tumors exhibited a distinct TME composition compared to ANXA4 high cases. ANXA4 low tumors showed higher levels of tumor infiltrating lymphocytes accompanied by increased deposition of acellular extracellular matrix. Further transcriptomic analysis demonstrated major alterations in transcriptional signatures related to epithelial-mesenchymal transition (EMT) and immune signaling. Transcription factor enrichment analysis and further functional validation identified ELF3 as one central regulator of invasive properties. Our integrative approach including molecular analyses with advanced histopathological segmentation uncovered novel roles for ANXA4 in modulating acute membrane repair, transcriptional regulation, and shaping cellular composition of the ccRCC tumor microenvironment.

## Introduction

Renal carcinoma affects worldwide more than 400.000 new patients per year, where clear cell renal cell carcinoma (ccRCC) is the most prevalent epithelial cancer subtype.^1^ While a variety of tumor driver genes have been identified (mostly related to epigenetic modulation such as *PBRM1*, *SETD2*), the common molecular underpinning is the early loss of *VHL* gene function (in sporadic as well as hereditary forms) translating into aberrant accumulation of HIF-1/-2 transcription factors resembling a pseudohypoxic state.^2,3^ HIF-1/-2 dependent transcriptional target genes (e.g. *VEGF*) drive the development of neovascularization within ccRCC tumors.^4,5^ Aside from this prominent vascularization, ccRCC is a prime example for the Warburg effect which is also evident at the morphological level since conventional FFPE (formalin-fixed and paraffin embedding) specimen preparation leads to exclusion of stored lipids and glycogen pools from individual cancer cells finally giving the impression of “clear cells” with prominent cellular boundaries.^6^

Prognosis, outcome and related clinical management of ccRCC is to date largely depending on respective tumor stage. With the advent of “omics” technologies (e.g. whole exome sequencing, transcriptomics, proteomics) and immune checkpoint blockade-based therapy protocols the role of molecular subclasses and the tumor (immune) microenvironment as well as the unmet need of precise biomarkers for these became evident.^7–9^ To develop more precise molecular classifications, previous studies identified favorable *PBRM1*-mutated subtypes in contrast to cases with more aggressive *BAP1* mutations.^9^ While additional genomic biomarkers (e.g. ClearCode34, CCP Score) based on selected gene signatures have been developed to predict prognosis in ccRCC, broad clinical application is still not established.^10^ On the contrary, current standardized pathological assessment (based on WHO guidelines) of ccRCC also grades for certain morphological growth patterns such as rhabdoid or eosinophilic appearance, as these features correlate with increased aggressiveness and poor clinical outcome.^11–13^ ccRCC shows high levels of intratumor heterogeneity in terms of morphological features including pure solid tumor sheets, cystic degeneration, stroma-enrichment ranging to (at least partial) papillary growth patterns.^14^ Interestingly, the growth of ccRCC follows a rather expansive than invasive mode resulting in the formation of compression of adjacent renal parenchyma and formation of a pseudo-capsule at the tumor/normal adjacent tissue interface.^15^ This mode of expansion presumably translates into increased mechanical forces on growing tumor nodules and requires specialized as well as adaptive membrane composition in respective ccRCC cell populations.

This generated our interest in a prominent, membrane-associated protein family of annexins, which share a common ANXA core-unit mediating Ca^2+^ dependent membrane binding via phospholipid interactions.^16^ ANXA proteins show a tissue- and cell-type specific expression profile, where the majority of respective proteins is detectable in epithelial cells.^17^ Given their wide propensity to interact with phospholipids of cellular membranes (e.g. plasma membrane), ANXA proteins have been implicated in a wide variety of pathologies ranging from inflammation, vascular homeostasis, neurological diseases as well as cancer.^16^ Based on previous studies selected ANXA subtypes have been studied in the context of ccRCC, namely ANXA1 to ANXA5, ANXA8 and ANXA13.^18–22^ In most of aforementioned studies increased expression levels of annexins correlated with poor prognosis and progression of disease in affected patients. Mechanistic in vitro studies employing knockdown or overexpression approaches indicated a direct impact of most ANXA proteins on cellular features such as proliferation, migration or invasion. In this context, ANXA3 seems to act in differing biological pathways, as a previous report demonstrated decreased levels of ANXA3 in ccRCC and identified an involvement of ANXA3 in lipid storage of respective cancer cells.^20^ While most ANXA proteins appear to modulate similar cellular mechanisms these proteins do not only exhibit a heterogeneous tissue- and cell-expression pattern, but further modulate cellular features most likely also in a disease-specific context.^16^

Employing published proteomic datasets, we globally analyzed the annexin protein family in samples of ccRCC (covering sporadic as well as syndromic VHL-disease related cases). Across three independent data sets, we identified ANXA4 as one of the most significantly enriched proteins in ccRCC tumor cells. Previous studies have already reported increased protein levels of *ANXA4* within ccRCC tumor tissue. ^23^ However, the functional role of ANXA4 in ccRCC remains to be established. To comprehensively evaluate relevant cellular functions of ANXA4 in ccRCC (and in particular for ANXA4 in oncological diseases)^24^ we employed genetic titration studies (knockdown, CRISPR/Cas9 editing as well as overexpression) and combined these with analyses on the tissue level using machine-learning-based morphometric segmentation. Our observations in vitro and in situ imply that ANXA4 exhibits differing subcellular localization patterns (namely cellular and nuclear membranes), which relate to ANXA4-dependent acute plasma membrane repair as well as transcriptional alterations in ccRCC.

## Results

### ANXA4 shows high specificity for ccRCC tumor cells with nuclear and cytoplasmic subcellular localization patterns

For global annotation and analysis of annexin family proteins, we employed available and independent proteomic datasets of ccRCC tumor tissue (as well as corresponding transcriptomic data for the CPTAC dataset; including our own analysis on syndromic cases of ccRCC in the context of VHL-disease (dataset C).^25–27^ While the compared datasets differed in numbers of detected proteins and analyzed patient cases, a robust detection for the majority of annotated annexin proteins was observed (Figure 1a and Supplementary Spreadsheet S1). Of note, only ANXA8, ANXA8L as well as ANXA10 were not detected based on reported cut-off parameters. When comparing ccRCC tumor tissue to adjacent normal renal parenchyma, ANXA4 protein showed the highest level of protein abundance (Figure 1a). Due to the bulk nature of conventional proteomic tissue analysis, we aimed for a more granular dissection in terms of cellular distribution of ANXA4. For further delineation of cellular identities, we interrogated available scRNA-Seq on ccRCC tumor tissue via the CELL×GENE platform.^28,29^ Interestingly, certain annexin family members such as ANXA2, which showed significant levels of enrichment in the bulk proteomic datasets, exhibited almost equal expression levels in tumor and endothelial cells as detected via scRNA-Seq (Figure 1b and supplementary Figure 1a). In this context, ANXA4 showed the highest level of tumor cell specificity when directly compared to other annexin genes (Figure 1b&c). To further corroborate these observations derived from proteomic and scRNA-Seq studies on a (sub-) cellular level, we employed immunohistochemistry on ccRCC tumor tissue. In line with scRNA-Seq based cellular deconvolution, we detected tumor and endothelial cell positivity for ANXA2 and ANXA3, whereas ANXA4 presented with exclusive signals within tumor cells (Figure 1d-g). In contrast to ANXA2 (and to a lesser extent also ANXA3) showing only a prominent membranous localization pattern, ANXA4 was also detected in the nuclear and nuclear membrane compartment of respective tumor cells (Figure 1g). To estimate the potential influence of ANXA4 on relevant parameters in respective patient populations, we employed the ccRCC TCGA data resource for correlative analysis.^7^ Remarkably, respective patient populations with low levels of ANXA4 were attributed a rather aggressive disease course reflected by reduced overall survival (Figure 1h and Supplementary Spreadsheet S2), while no significant differences in tumor stage were identified (Figure 1i). In line with decreased survival parameters, higher tumor grades and increased levels of positive lymph nodes correlated with respective ANXA4 low tumors (Figure 1j&k). Together, our analysis demonstrated a selective expression pattern of ANXA4 in ccRCC tumor cells with various subcellular localization patterns (membranous and nuclear) and indicated a correlation of low ANXA4 expression with a more aggressive disease course.

**Figure 1.**
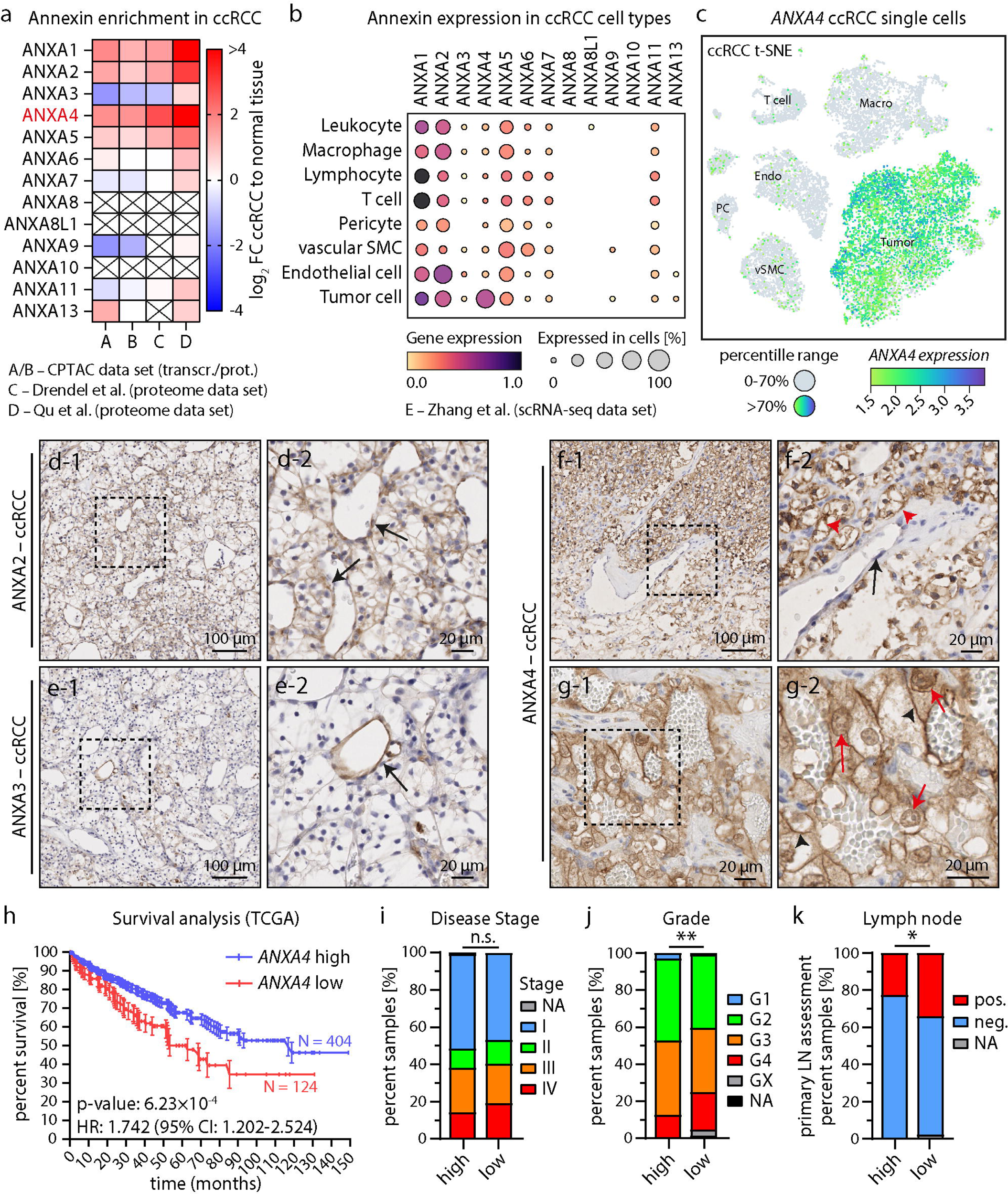
ANXA4 is upregulated in ccRCC, prognostic favorable and recruited to cytoplasm and nuclear membranes. (a) Heatmap depicts the expression of annexin family members in previously published transcriptome and proteome datasets of indicated ccRCC cohorts compared to normal adjacent tissue (FC – fold change; crossed out boxes – not detected).^25–27^ (b) Dot plot shows relative expression of annexin family genes in indicated cell populations derived from previously published scRNA-Seq analysis of seven ccRCCs.^28^ (c) t-SNE blot indicates expression of *ANXA4* in single cell populations in ccRCC (cells below the 70-100% percentile range of continuous values for *ANXA4* expression are gray labeled). (d-g) Immunohistochemistry analysis of ANXA2, ANXA3 and ANXA4 demonstrated differential expression of annexin proteins (dashed boxes indicate regions of detail images). ANXA2 localized to endothelial (black arrows) and cancer cells, ANXA3 selectively to endothelial cells and ANXA4 selectively to cancer cells (red arrowheads). Detailed analysis of subcellular localization (g) of ANXA4 revealed distinct enrichment at the cytoplasm (back arrowheads) and nuclear membrane (red arrows) of ccRCC cells. (h) Survival analysis for *ANXA4* comparing patients with high (N=404; blue) and low (N=124; red) expression in the ccRCC TGCA cohort (HR – hazard ratio of *ANXA4* low to high expression; CI – ±95% confidence interval). (i-j) Comparison of *ANXA4* high (N=404) and low (N=124) expressing ccRCCs in the TGCA cohort for disease stage, histologic grade and primary lymph node presentation (n.s. – not significant; * p<0.05; ** p<0.01).

### Loss of VHL contributes to upregulation of ANXA4 renal cancer cell lines

Given the pronounced expression pattern of ANXA4 in ccRCC tumor cells in situ, we searched for appropriate cellular model systems to delineate functional implications of ANXA4. Here, we globally analyzed renal carcinoma derived cell lines in the Cancer Cell Line Encyclopedia and mapped expression levels of annexin proteins to *VHL*-mutations or loss of the p-arm of chromosome 3 (Figure 2a, Supplementary Figure S1 and Supplementary Spreadsheet S1).^30^ Loss of chromosome 3p (encompassing ccRCC driver genes like *VHL*, *PBRM1*, *SETD2*, *BAP1*) or mutations of the *VHL* gene are pathognomonic features of ccRCC.^3^ In line with our tissue-based analysis (see Figure 1), *ANXA4* showed again a consistent elevation of expression levels in numerous kidney tumor cell lines. Correlation to *VHL*-mutation status and loss of chromosome 3p indicated a potential link between *ANXA4* and *VHL*-expression levels. To further corroborate these observations, we made use of two independent ccRCC cancer cell lines with natural pVHL-loss (A498 and 786-O). Proteomic analysis of respective cell lines with pVHL-loss or constitutive pVHL-re-expression demonstrated a pronounced dependence of ANXA4 protein levels on the presence of pVHL (Figure 2b – most prominently in the A498 cell line and Supplementary Spreadsheet S3). For further functional analysis, we generated knockdown lines on these cellular backgrounds and respective overexpression on a non-cancerous renal proximal tubular epithelial cell (RPTEC) line (Figure 2c-e and Supplementary Figure S1). Previous studies have reported on ANXA4 affecting cellular features such as proliferation.^31,32^ To investigate if ANXA4 might impact on ccRCC proliferative behavior, we employed respective cellular models and recorded growth dynamics using long-time live imaging (up to 120h). In contrast to previous reports,^32,33^ we did not observe any significant impact of ANXA4 protein on cellular proliferation neither in knockdown nor in overexpression conditions (Figure 1f-h). To exclude insufficient knockdown efficiency, we further made use of CRISPR/Cas9 genome editing and generated complete *ANXA4* knockout clones from two independent guide RNAs on the A498 cellular background (given the most prominent dependency on pVHL-protein – Figure 2i&j). Of note, the knockout of *ANXA4* did not translate into compensatory upregulation of other annexin proteins such ANXA2 or ANXA3 (Figure 2j). Interestingly, also the complete knockout of *ANXA4* in A498 cells did not affect cellular proliferation as measured via BrdU incorporation studies (Figure 2k). In summary, our studies revealed (at least partial) pVHL-dependency of *ANXA4* gene products in ccRCC cancer cell lines and further demonstrated that ANXA4 protein levels do not modulate cellular proliferation.

**Figure 2.**
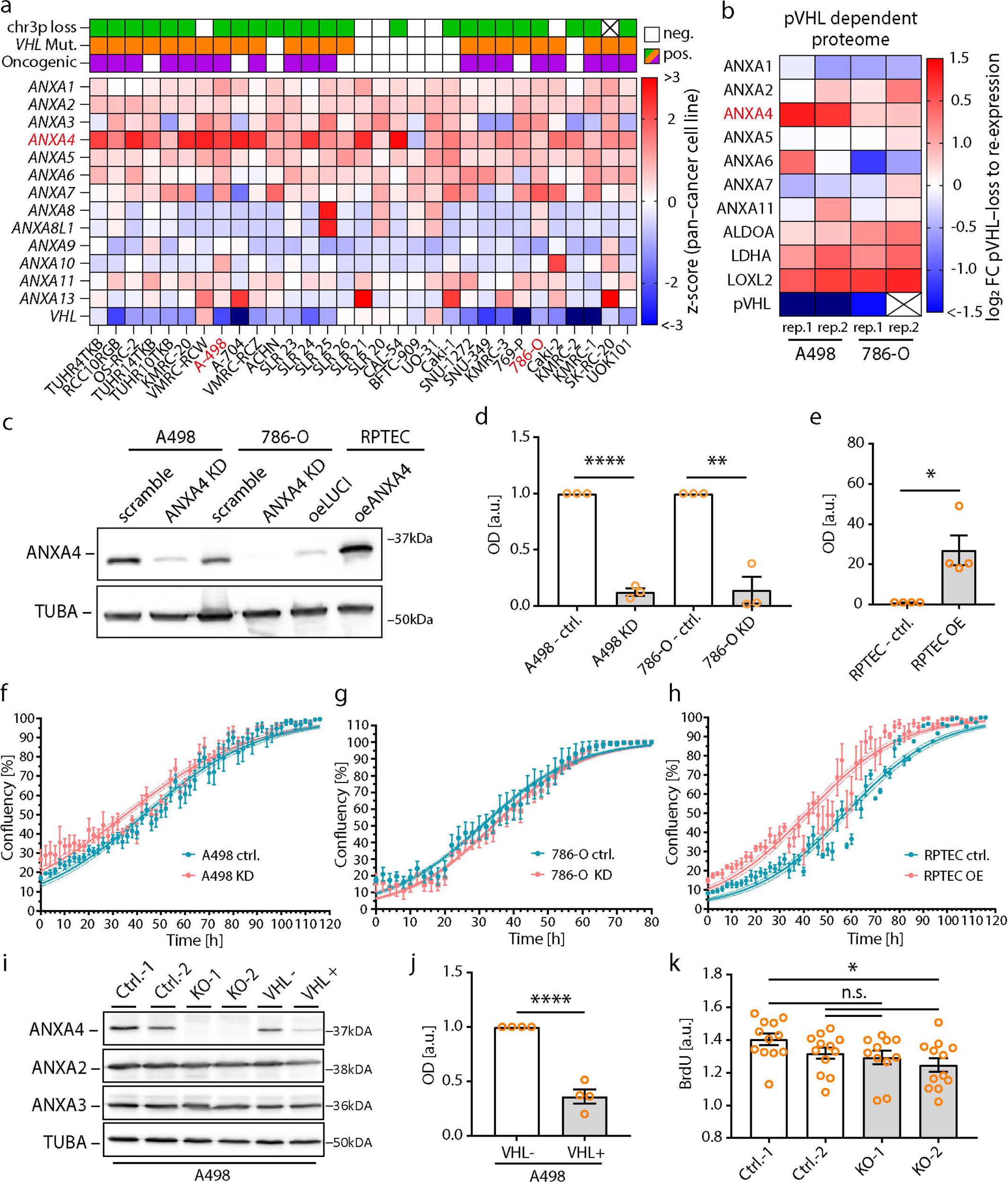
ANXA4 is upregulated in kidney cancer cell lines by loss of VHL. (a) Heatmap maps present the expression of *VHL* and annexin family genes in kidney cancer derived cell lines included into the Cancer Cell Line Encyclopedia database. Z-scores indicate gene expressions relative to the entire cancer cell line database as reverence. Estimated loss of the p-arm of chromosome 3 (chr3p), mutations of the *VHL* gene and prediction of oncogenicity (OncoKB) are indicated (colored box – positive, white box – negative for these features; crossed out box – no data available). A498 and 786-O cell lines (red label) were selected for further analysis of *ANXA4*. (b) Proteome analysis of A498 and 786-O ctrl. and pVHL re-expression cell lines demonstrated regulation of ANXA4 in a pVHL dependent manner (two replicates per cell line were analyzed). Heatmap shows relative log_2_ fold changes (FC) of pVHL-loss over re-expression for annexin family proteins and marker proteins for pVHL dependent regulation (crossed out box – no ratio could be calculated). (c-e) Western blot analysis for ANXA4 in shRNA mediated A498 *ANXA4* knockdown (KD) and 786-O *ANXA4* KD or overexpression (oeANXA4) in RPTEC cell lines (TUBA was used as loading control). Scramble shRNAs or Luciferase (oeLUCI) were expressed as controls (ctrl.). Densitometry of western blots shows optical densities of three (d) or four (e) independent experiments. Arbitrary units (a.u.) were normalized to respective controls per experiment. (f-h) Analysis of cell growth curves revealed no differences in *ANXA4* KD cell lines and only slight differences in ANXA4 overexpressing (OE) RPTECs (one representative experiment out of several independent experiments is shown; dots indicate mean values and error bars S.E.M. per time point; growth curves indicate fitted growth rates (nonlinear regression) and dotted line indicates ±95% confidence intervals). (i&j) Western blot analysis for ANXA4 in *ANXA4* knockout (KO) and pVHL re-expression A498 cells (ANXA2, ANXA3 and TUBA were used as reference). CRISPR/Cas9 generated ctrl.-1&-2 and KO-1&-2 indicate monoclonal cell lines using independent non-targeting or *ANXA4* specific guide RNAs per cell line. Densitometry of western blots confirmed downregulation of ANXA4 following pVHL re-expression (VHL+) in A498 cells (four independent replicates normalized to respective controls per experiment were analyzed). (k) Cell proliferation analysis by BrdU incorporation in *ANXA4* KO and ctrl. cell lines (12 replicates per genotype out of 4 independent experiments and 3 replicates per experiment; a.u. – arbitrary units).Dot plots show analyzed replicates (error bars indicate mean and S.E.M.; n.s. – not significant; * p<0.05; ** p<0.01 and **** <0.0001).

### Membrane stress regulates subcellular localization of ANXA4 in ccRCC cancer cells

Based on our observations of distinct localization patterns for ANXA4 in ccRCC in situ (Figure 1), we further characterized subcellular distribution of ANXA4 in cellular models of renal carcinoma. While ANXA4 presented with a predominant membranous localization in situ, ANXA4 showed a prominent cytoplasmic pattern under steady state condition in vitro (Figure 3a). As previous studies have reported about the potential involvement of annexin proteins in processes of membrane repair, we exposed the A498 carcinoma cell line to stress conditions such as hypoosmolarity as well as digitonin permeabilization.^34^ In both conditions, a rapid recruitment of ANXA4 towards the cellular membrane as well as a very distinct accumulation at the nuclear membrane became evident (Figure 3b&c). Interestingly, also cellular growth conditions appeared to modulate localization of ANXA4 as high cellular densities led to an almost complete exclusion from the nuclear/nuclear membrane compartment (Supplementary Figure S2). Based on the rapid recruitment of ANXA4 towards membrane structures upon acute stress exposure, we tested the impact of ANXA4 on processes such as cellular membrane repair employing short-term digitonin treatment (Figure 2g). In fact, lower levels of ANXA4 resulted in decreased capacity for membrane repair and higher numbers of disrupted cells (Figure 3h&i). As these observations indicate a dynamic and context dependent mode of localization for ANXA4, we further aimed to characterize distribution patterns in patient samples with differing biological behavior. Given its prominent role of plasma membrane integrity in cancer cell invasion, we analyzed localization of ANXA4 employing immunohistochemistry on a cohort of ccRCC cases with non-progressive versus progressive disease (Supplementary Spreadsheet S4).^35^ Remarkably, we observed a predominant membranous pattern of ANXA4 in progressive cases when compared to non-progressive cases of ccRCC, whereas the latter showed a more frequent localization at the nuclear/nuclear membrane compartment (Figure 3j&k). Altogether, these experiments demonstrate the versatile subcellular localization patterns of ANXA4 in ccRCC, which are presumably linked to the involvement of ANXA4 in processes of acute membrane repair.

**Figure 3.**
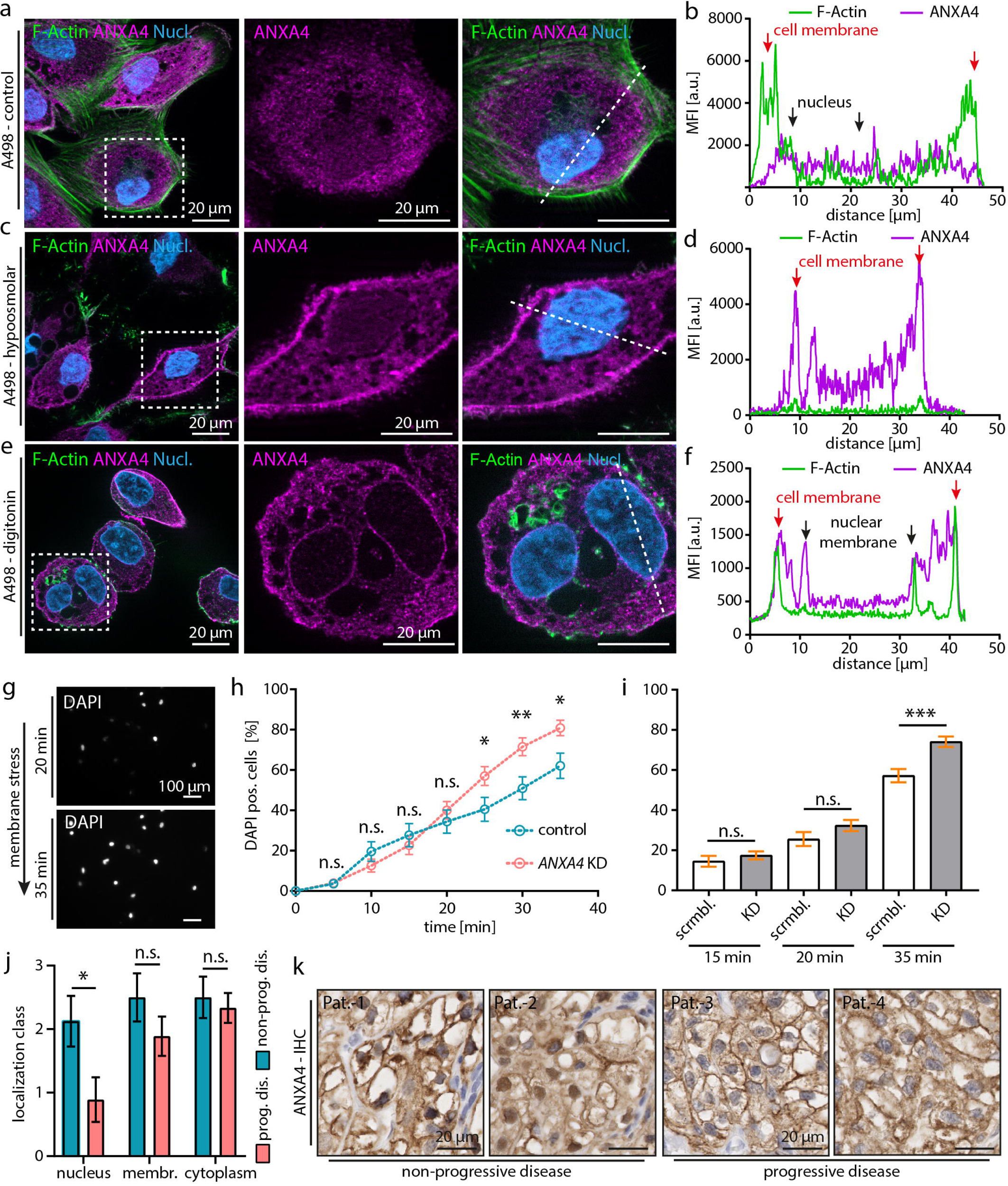
Membrane stress regulates subcellular localization of ANXA4. (a-f) Membrane stress by hypoosmolar media or digitonin treatment induced recruitment of ANXA4 to the nuclear membrane in A498 cells. Representative immunofluorescence images show subcellular localization of ANXA4 (cells were co-stained by Phalloidin (F-Actin) and Hoechst (Nucleus)). Dashed boxes indicate regions of magnification. Dashed lines indicate positions of correlating line scans. Line scans (b&d&f) show mean fluorescence intensities (MFI) as arbitrary units (a.u.) of ANXA4 and F-Actin (red arrows indicate cytoplasm membranes and black arrows indicate nuclear membranes). (g) Representative fluorescence images show uptake and nuclear staining of DAPI in life cells over time after induction of membrane damage by digitonin. (h) Quantification of one representative experiment for digitonin induced DAPI uptake over 35 minutes (*ANXA4* KD and control A498 cells; dots indicate mean and S.E.M of 12 analyzed region of interests per genotype). (i) Quantification of digitonin induced DAPI uptake into *ANXA4* KD and control A498 cells for indicated time points (N=12 regions of interest per genotype, time point and experiment and 3 independent experiments (total N=36) were analyzed). (j&k) ANXA4 localization analysis in ccRCC patients with progressive (N=9) or non-progressive (N=9) disease. Localization of ANXA4 was quantified into three classes – cytoplasm, nucleus and plasma membrane (a four tier score was applied: 0 – no staining to 3 strong staining). Representative immunohistochemistry (IHC) images showing subcellular localization of ANXA4 (brown) in ccRCCs of four representative patients. Nuclei were stained by hematoxylin (blue). All error bars indicate mean and S.E.M.; n.s. – not significant; * p<0.05; ** p<0.01; *** p<0.001.

### ANXA4 correlates to features of the tumor microenvironment of ccRCC

Our initial characterization of the annexin protein family identified ANXA4 as one of the most selectively expressed annexins in ccRCC tumors (see Figure 1). Interestingly, complementary in vitro studies did not reveal a major impact on tumor inherent features such as proliferation; whereas population-based analysis indicated that lower levels of ANXA4 correlate with decreased overall survival of affected patients (see Figure 1-3). We therefore aimed to characterize the composition of ccRCCs (cellular and non-cellular) on the tissue level in correlation to ANXA4 abundance to identify relevant features potentially influencing tumor biology. Here, we made use of comprehensively characterized ccRCC cases including genetic, transcriptomic, proteomic and histology data (i.e. “Clinical Proteomic Tumor Analysis Consortium – CPTAC”).^8,27^ Based on mRNA expression and proteome abundance of ANXA4 we selected for ANXA4-high and -low cases (Figure 4a and Supplementary Spreadsheet S5), and subsequently annotated cellular composition of respective tumor samples employing machine learning based identification and segmentation on a single cell level. Using this detailed topographical annotation, we performed proximity and cellular neighborhood analysis, which allowed for regional clustering of cellular neighborhoods and pattern analysis (Figure 4b&c). Compositional analysis of these individual regions indicated that lower levels of ANXA4 correlated to increased levels of lymphocyte presence within respective tumor samples (Figure 4c and Supplementary Spreadsheet S5). To validate this global neighborhood analysis, we extracted granular cellular segmentation and quantification confirming a higher presence of infiltrating lymphocytes in ANXA4-low tumors (Figure 4e&f). These observations were further substantiated by significantly different levels of lymphocyte clustering and lymphocyte-tumor cell proximities (Figure 4g-l), as well as correlation of ANXA4 levels to global immune scores in the ccRCC CPTAC cohort (4m&n). Further deconvolution of immune cell subtypes indicated an increase of lymphocyte (i.e. CD3, CD4, CD8, DC19, CD20, CD69, CD103), monocyte/macrophage (i.e. CD14, CD33, CD68), dendritic cell (i.e. CD11) and granulocyte cell (i.e. CD66b) populations in ANXA4-low tumor samples (Figure 4o). As the global regional analysis also highlighted differences in the stroma compartment (Figure 4b), we further analyzed stroma and stroma cellularity in respective ANXA4-high and -low tumors. Remarkably, number and distribution of individual stroma cells did not drastically differ between both groups, whereas areas of deposited stroma (non-cellular) were significantly increased in ANXA4-low cases (Figure 5a-j and Supplementary Spreadsheet S5). These findings indicate that ANXA4 abundance correlates with deposition of extracellular matrix (ECM) within the tumor microenvironment of ccRCC. Further mapping of relevant ECM components demonstrated indeed higher levels of collagens and laminins in respective ANXA4-low tumor samples (Figure 5k). Thus, our results show that ANXA4 levels correlate to distinct features of the tumor microenvironment of ccRCC characterized by differing lymphocyte composition as well as stroma/ECM deposition.

**Figure 4.**
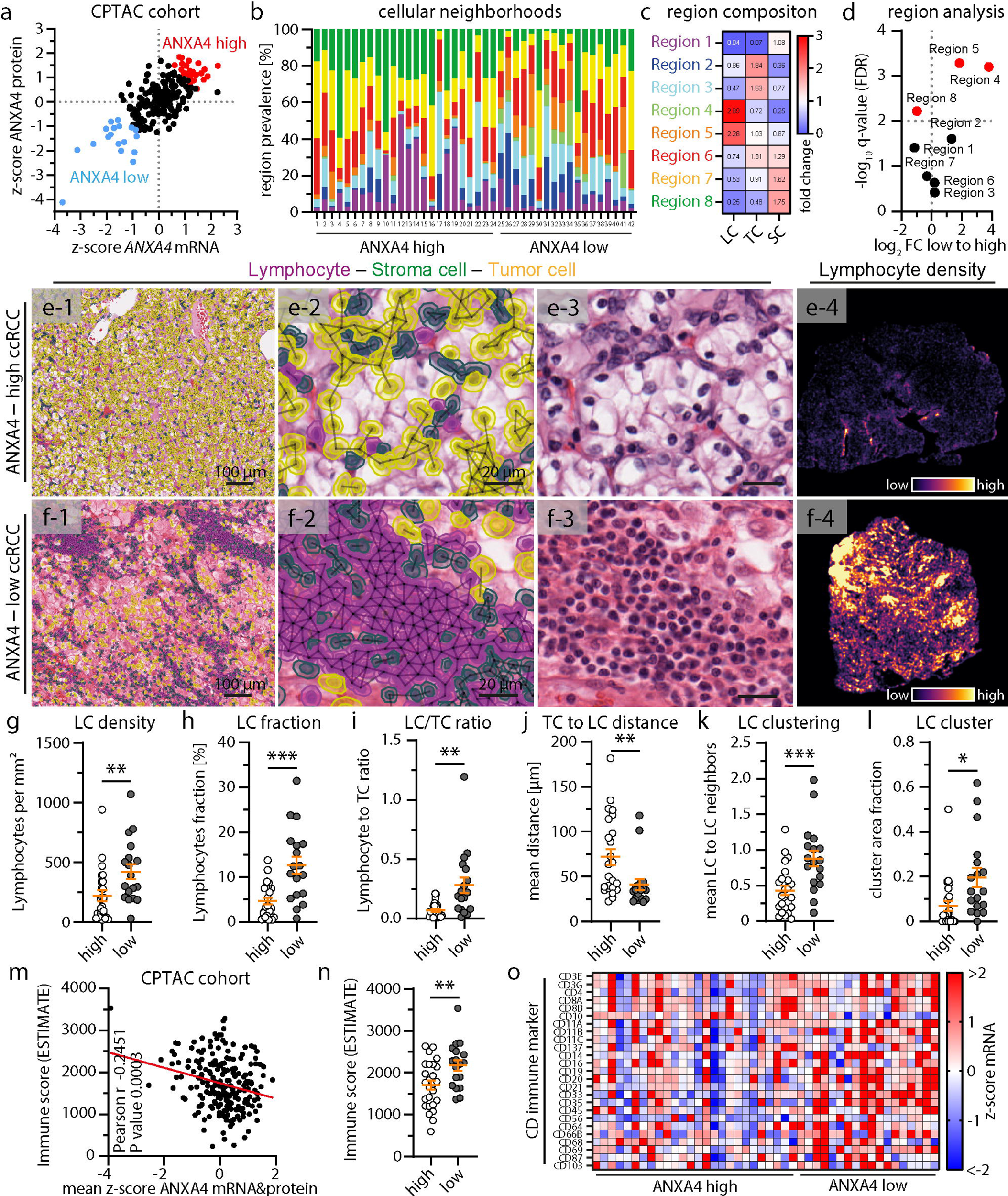
ANXA4 modulates cellular neighbourhoods and lymphocyte infiltration in ccRCC. (a) ANXA4 expression analysis on mRNA and protein level identified ANXA4 low (blue) and high (red) cancer samples in the ccRCC CPTAC cohort (N=213). (b-d) Analysis of ANXA4 high (N=24) and low (N=18) ccRCCs for cell neighborhoods shows eight distinct tissue regions (neighborhood types) defined by cell type composition and density. Dot plot graph shows fold changes (FC) in region prevalence and reveals significant higher prevalence of Lymphocyte (LC) enriched regions in ANXA4 low cancer samples (TC – tumor cells, SC – stroma cells; red dots indicate significant regulated regions defined as FDR q-value <0.01). Color-code for regions and region composition in shown in heatmap (c). Region composition of ANXA4 high and low ccRCC samples is depicted as stacked bar chart in (b). (e-f) Representative images showing machine learning based cell segmentation and classification in HE stained tissue section from the CPTAC cohort (connecting lines indicate neighboring cells within 15 µm; colors of cell overlays label cell types as indicated). Section overviews (e-4&f-4) show heatmaps for lymphocyte density of representative ANXA4 high and low cancer samples (low – black to high – yellow density as indicated). (g-l) Quantitative analysis shows increased lymphocyte density (g), percentage (h) and lymphocyte to tumor cell ratios (i) in ANXA4 low ccRCCs. Tumor cell distance to the next adjacent lymphocyte was reduced (k) and lymphocyte cluster formation was increased (k&l), as indicated by analysis of neighboring lymphocytes and area fraction of lymphocyte clusters. (m&n) Correlation analysis of ANXA4 expression and ESTIMATE immune scores demonstrated significant negative correlation in the CPTAC cohort (N=213) (m) as well as increased immune scores in the ANXA4 low group (n). (o) Heatmap of z-scores of selected cluster of differentiation (CD) markers for immune cells for ANXA4 high and ANXA4 low samples in the CPTAC transcriptome dataset (z-scores are relative to all cancer samples in the dataset). Dot plots show analyzed ANXA4 high (N=24) and ANXA4 low (N=18) patients (error bars indicate mean and S.E.M.; * p<0.05; ** p<0.01; *** p<0.001 and **** <0.0001).

**Figure 5.**
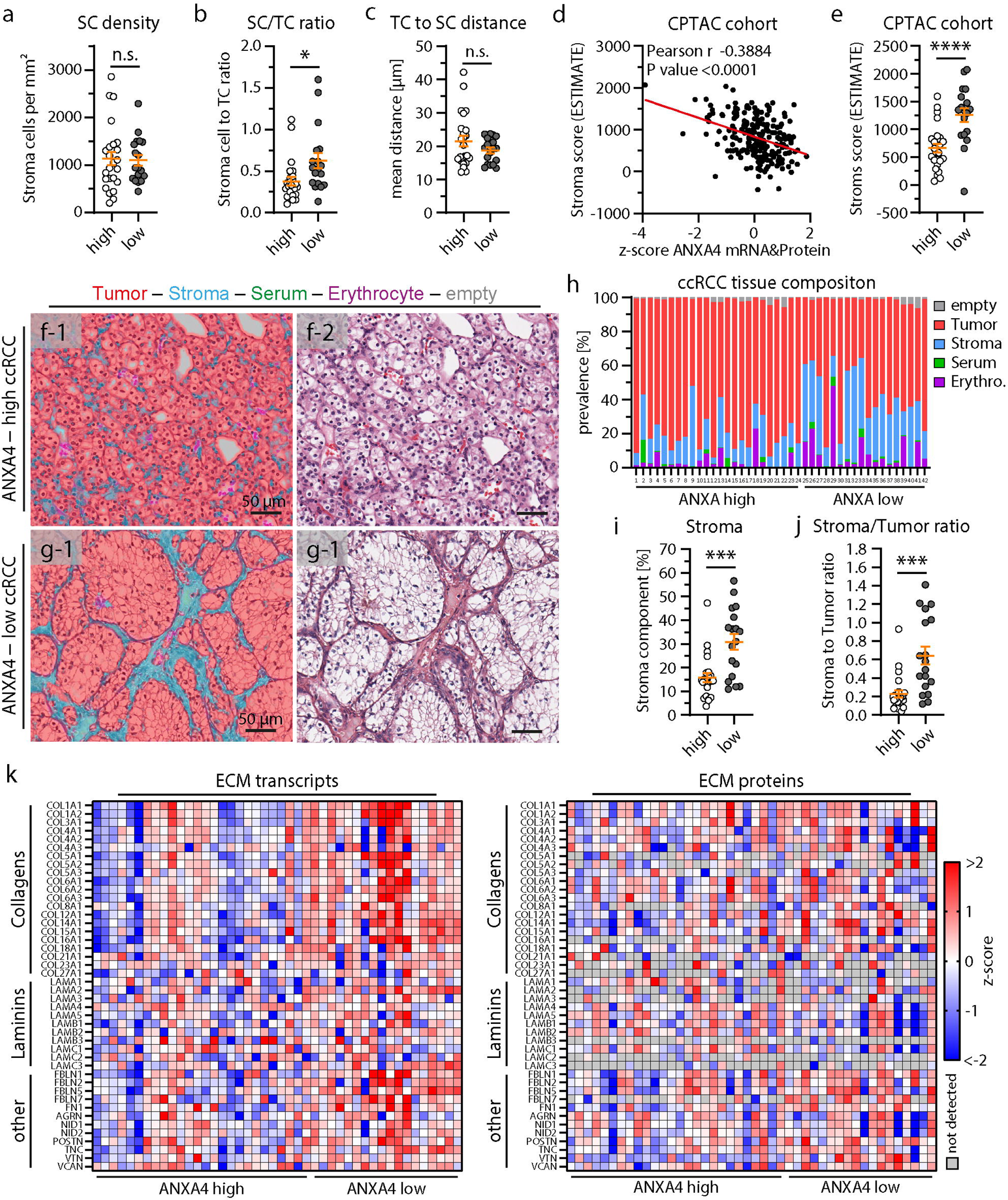
ANXA4 expression correlates with stroma content in ccRCCs. (a-c) Analysis of stroma cell (SC) density (a), stroma cell to tumor cell (TC) ratio (b) and tumor cell distance to the next adjacent stroma cell (c) indicates only mild alterations comparing ANXA4 high and low ccRCCs. (d&e) Correlation analysis of ANXA4 expression and ESTIMATE stroma scores demonstrated significant negative correlation in the CPTAC cohort (N=213) as well as increased stroma scores in the ANXA4 low group. (f&h) Representative images depict machine learning based annotation of tissue types in HE stained tissue section from the CPTAC cohort. Tissue classes were annotated as tumor tissue, stroma tissue, erythrocyte filled space, serum filled space or empty space. Image overlays (f-1&g-1) indicate detection probabilities for indicated tissue types/classes (high color intensity indicates high detection probability for respective classes). Tissue composition of ANXA4 high and low ccRCC samples is shown as a stacked bar chart in (h). (i&j) Analysis of stroma tissue prevalence (i) and stroma to tumor tissue ratio (j) revealed an increase of the stroma component in the ANXA4 low ccRCC group. (k) Heatmap of z-scores of selected ECM gene products in ANXA4 high and ANXA4 low samples in the CPTAC transcriptome and proteome dataset (z-scores are relative to all cancer samples in the dataset). Dot plots show analyzed ANXA4 high (N=24) and ANXA4 low (N=18) patients (error bars indicate mean and S.E.M.; n.s. – not significant; * p<0.05; *** p<0.001 and **** <0.0001).

### Loss of ANXA4 influences transcriptional signatures of ccRCC cancer cells

Subcellular localization studies in vitro and in vivo (see Figure 1 and Figure 3) demonstrated distinct and dynamic recruitment properties of ANXA4 towards the cellular plasma membrane as well as the nuclear compartment. In addition, previous studies demonstrated a potential interaction of NF-kappa-B and recruitment of ANXA4 towards the nucleus, thereby influencing transcriptional activity.^36^ To further dissect the potential modulating impact of ANXA4 on transcription, we performed mRNA-sequencing studies using our *ANXA4* knockout model of A498 cells (Supplementary Spreadsheet S6). Principal component analysis showed a clear separation of knockout and respective control clones, while in addition also the different clonal background was apparent (Figure 6a). Differential expression analysis and unsupervised clustering identified 834 significantly regulated genes between both conditions (Figure 6b&c). Of note, a compensatory upregulation of other annexin family genes was not detected in the respective *ANXA4* knockout condition. Interestingly, several genes/transcripts related to regulation or composition of the ECM (such as *LOXL3* or *TNC*) were significantly downregulated in conditions of ANXA4 loss (Figure 6b). On the contrary, distinct upregulation of metabolic enzymes (e.g. *HSD3B7*), immune regulatory related genes (e.g. *NFKBIA*, *IL34*, *IL7R*) as well as transcriptional regulation (e.g. *ELF3*, *RASSF7*) was observed in respective ANXA4 knockout clones (Figure 6b). Global analysis employing gene ontology-based (GO) annotations further demonstrated the impact of ANXA4 on extracellular matrix structure and turnover, cell membrane dynamics, as well as regulation of immune-related GO-terms such as T-cell mediated cytotoxicity or inflammatory response (Figure 6d&e and Supplementary Spreadsheet S6). These transcriptional alterations might therefore indirectly reflect our observations of differential cellular compositions in terms of immune cell infiltrates or stromal structure in correlation to ANXA4 protein abundance (see Figure 4&5). To distill underlying transcriptional regulation, we performed intersectional analysis of most significantly regulated genes with databases comprising analyzed biological concepts (e.g. Matrisome, EMTome – epithelial mesenchymal transition), which led to a core set of 38 genes with the majority of genes related to epithelial-mesenchymal-transition (Figure 6f and Supplementary Spreadsheet S6). Further transcription factor enrichment analysis revealed the ETS-family member *ELF3* as the most significantly enriched transcription factor of upregulated genes in ANXA4 loss conditions (Figure 6g and Supplementary Spreadsheet S6). E74-like factor 3 (ELF3) is involved in various cellular processes ranging from epithelial differentiation to modulation of cytokines, and has been shown to exert context-dependent roles either as a tumor-suppressor or even more oncogene-like functions in various malignancies.^37^ Taken together, our observations imply ANXA4-dependent altered transcriptional signatures involving matrisomal- and immune-related genes, and led to the identification of the transcription factor *ELF3* as a potential regulator of relevant gene sets.

**Figure 6.**
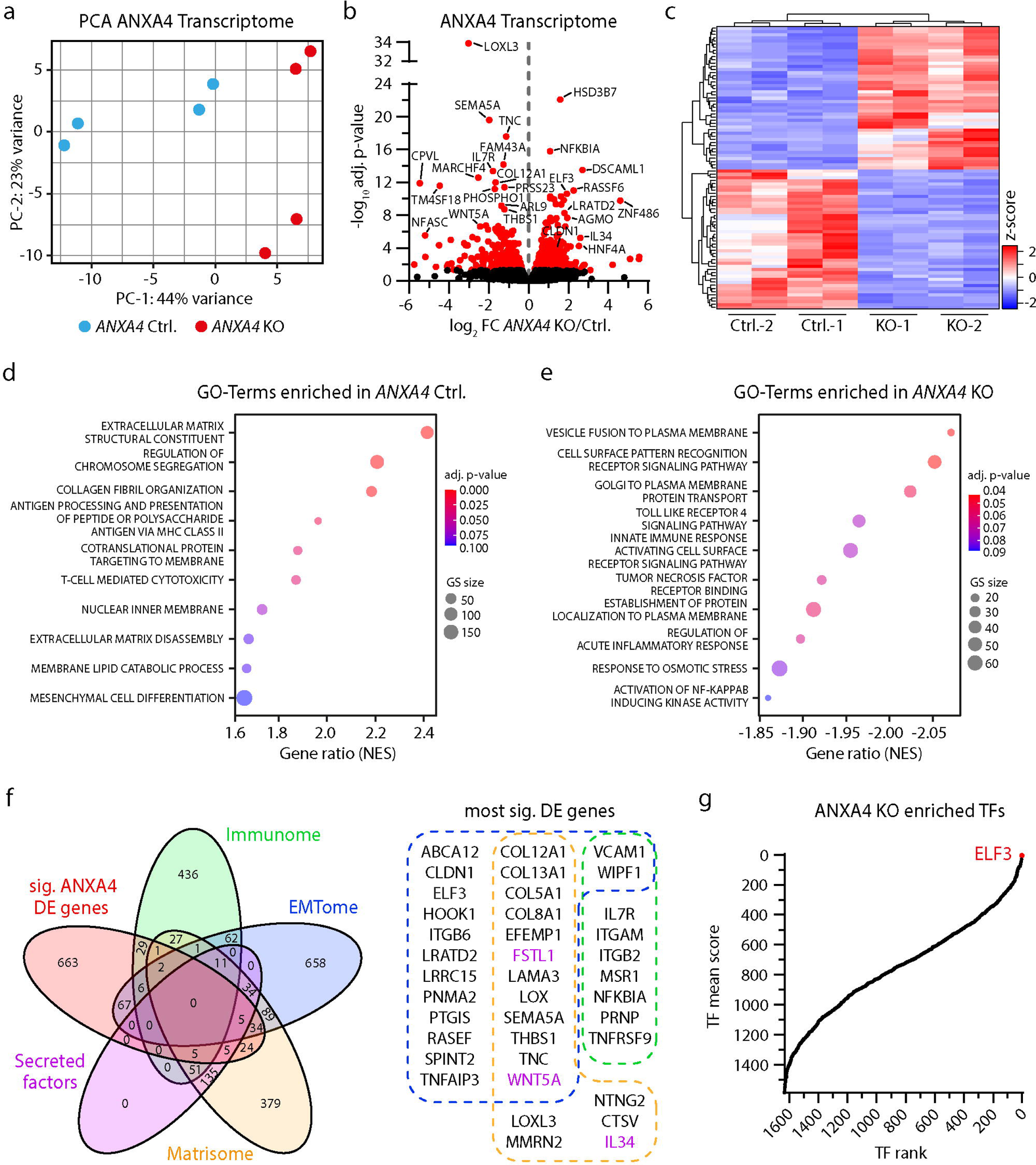
Transcriptome analysis of *ANXA4* KO cells reveals altered matrisome and immune signaling. (a) Principle component analysis (PCA) of transcriptome analysis (mRNA sequencing) of *ANXA4* Ctrl. and KO A498 cell lines. Two replicates of Ctrl.-1&-2 (blue) and KO-1&-2 (red) CRISPR/Cas9 clones were analyzed. (b) Volcano plot of differential expression analysis shows 834 significantly regulated genes (adjusted p-value <0.05; red dots; FC – fold change). Highly altered genes suggest regulation of extracellular matrix (e.g. *LOXL2*, *COL12A1*, *TNC*), immune signaling (e.g. *NFKBIA*, *IL34*, *IL7R*) and epithelial–mesenchymal transition (e.g. *ELF3*, *CLDN1*, *WNT5A*) related mechanisms. (c) Heatmap and clustering of z-scores of the most significant regulated genes (adjusted p-value <0.0001 and absolute log_2_ fold change >1) demonstrated robust regulation of these genes in all monoclonal cell lines and replicates. (d&e) Gene set enrichment analysis shows significant regulation (adj. p-value <0.01) of extracellular matrix, immune signaling, epithelial–mesenchymal transition, chromosomal regulation and nuclear membrane associated GO-Terms (NES – normalized enrichment score of KO to WT; GS – gene set size). (f) Venn-diagram of 834 significant differentially expressed (DE) genes by loss of ANXA4 and annotation of the transcriptome to databases for immune (Immunome), extracellular matrix (Matrisome, including secreted factors) and epithelial–mesenchymal transition (EMTome) related genes. The Gene list shows the most significant DE genes (adjusted p-value <0.0001 and absolute log_2_ fold change >1) annotated to at least one database (colored boxes and gene symbols indicate annotation of genes to respective databases). (g) Transcription factor (TF) enrichment analysis indicates ELF3 as relevant for ANXA4-dependent regulated genes/gene sets.

### ELF3 suppresses invasive and migratory phenotypes

To further validate our transcriptome-based analysis (see Figure 6), we evaluated ELF3 protein levels in respective *ANXA4* knockout cells and controls. Here, we not only confirmed increased levels of ELF3, but further demonstrated upregulation of CLDN1 in conditions of ANXA4 loss (Figure 7a-c and Figure 6b). Of note, albeit slightly differing detection levels for CLDN1 between individual knockout clones (Figure 7c), the increase of ELF3 was robustly detected (Figure 7b). Our initial analysis of *ANXA4* knockout cells did not reveal a pronounced impact on tumor inherent features such as proliferation (see Figure 2). However, we observed a relevant role for ANXA4 in mediating membrane repair upon cellular stress exposure (Figure 3). Invasive growth (related to more aggressive tumor biology) requires cancer cells to remodel their cellular morphology for adaptation towards environmental constraints such as matrix stiffness and extracellular pore sizes.^38^ Therefore, we hypothesized that a loss of ANXA4 translating into diminished membrane repair capacity might also affect invasive migration modes of respective cancer cells in vitro. To test this hypothesis, we challenged *ANXA4* knockout and control cells in conventional transwell migration assays but did not observe any significant differences (Figure 7d). However, when we further modified this experimental setup by adding extracellular matrix (Matrigel layer), *ANXA4* knockout cells did show diminished invasive migration compared to respective controls indicating that additional features beyond membrane repair influence this phenotype (Figure 7e&f). Given our transcription factor enrichment analysis (see Figure 6), we reasoned that ELF3 might be (at least partially) involved in the observed migratory phenotype. Therefore, we generated stable overexpression lines of ELF3 on a wild type *ANXA4* A498 cancer cell background line (Figure 7g). Remarkably, ELF3 overexpression not only translated into a phenocopy of impaired invasive migration capacity in *ANXA4* knockout cells (Figure 7h&i), but also impaired 2D migration of A498 cancer cells (Figure 7j&k). Together, these observations imply that ELF3 suppresses invasive migratory modes in line with its known role as a potential tumor suppressor and counter-acting transcription factor of EMT.^37^ Further co-expression analysis of *ELF3* employing the TCGA ccRCC database revealed *ELF3* co-regulated gene signatures (intersectional analysis with Immunome, EMTome and Matrisome), where secreted chemokine factors (e.g. CCL2, CXCL1, CXCL6) were identified (Figure 7l and Supplementary Spreadsheet S7). These observations suggest that ELF3 influences not only cellular phenotypes (e.g. migration, invasion) in a context-dependent manner, but also might modulate intercellular communication via secreted factors thereby shaping compositional features of the TME downstream of ANXA4.

**Figure 7.**
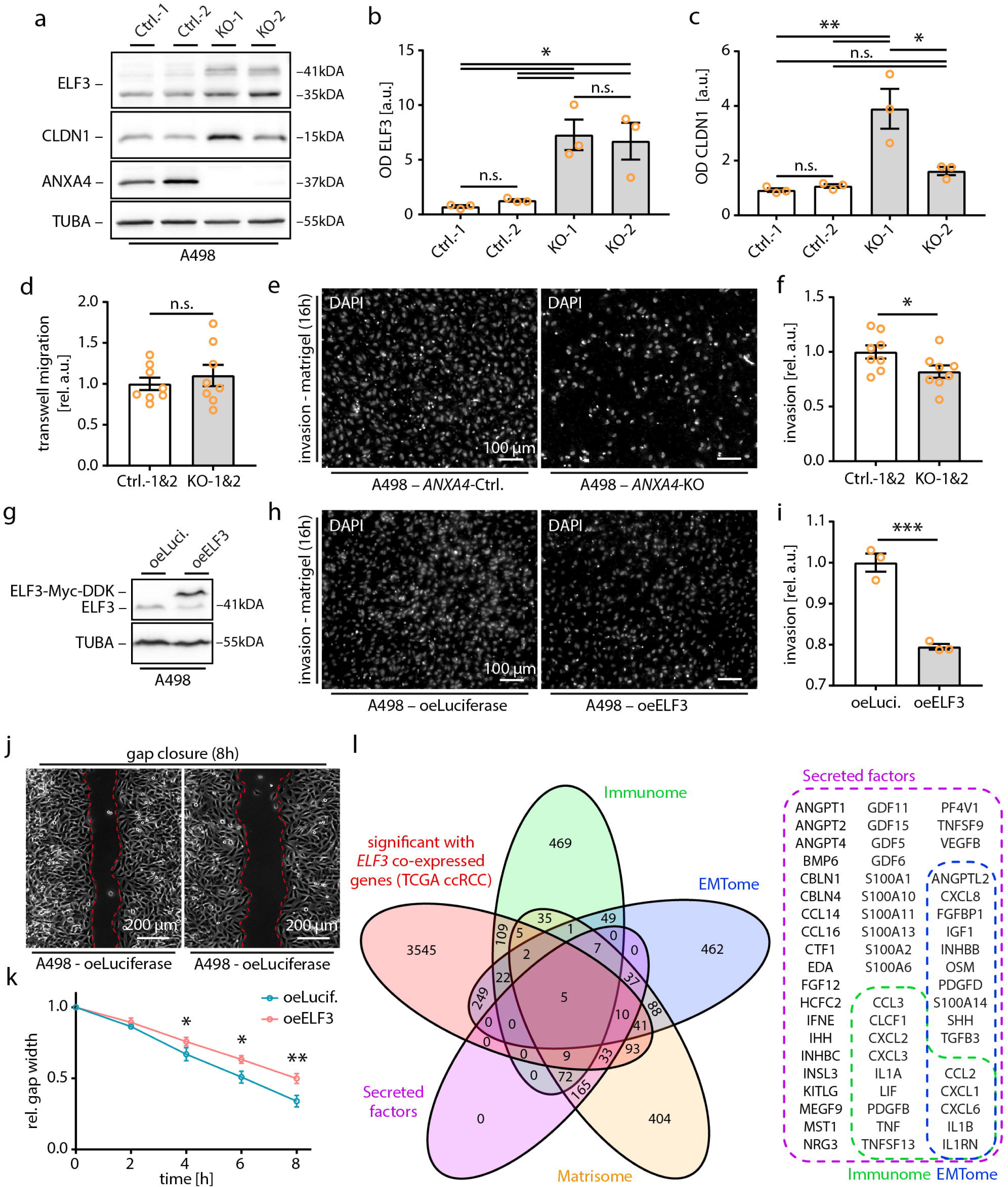
ELF3 mediates ANXA4 associated phenotypes in ccRCC. (a-c) Western blot analysis for ELF3 and CLDN1 confirms upregulation of ELF3 and CLDN1 in *ANXA4* KO cells (ANXA4 and TUBA are shown as reference). Densitometry of western blots shows optical densities (OD) as arbitrary units (a.u.). Three independent experiments were analyzed and ODs were normalized to the mean OD of Ctrl-1&-2 per experiment. (d) Analysis of transwell migration of *ANXA4* ctrl. and KO cells (Ctrl.-1&-2 and KO-1&2 cells were combined for statistical analysis; three replicates per genotype of three independent experiments were analyzed; a.u. – arbitrary units relative to control). (e&f) Analysis of cell invasion of *ANXA4* ctrl. and KO cells (Ctrl.-1&-2 and KO-1&2 cells were combined for statistical analysis; four replicates per genotype out of four independent experiments were analyzed). Representative fluorescence images of DAPI stained cell nuclei show transmigrated (invasive) cells after 16 hours. (g) Western blot analysis for ELF3 in *ELF3* overexpression (oeELF3; Myc-DDK tagged) and control (oeLuci – Luciferase overexpression) A498 cells (TUBA was used as reference). (h&i) Analysis of cell invasion of Luciferase (control) and ELF3 overexpression A498 cells (n=3). Representative fluorescence images of DAPI stained cell nuclei show transmigrated (invasive) cells after 16 hours. (j&k) Gap closure assay for Luciferase (control) and ELF3 overexpression (oeELF3) A498 cells. Representative phase contrast images show gap width after 8 hours. Quantification shows mean values (dots) of three independent experiments at indicated time points (gap width was normalized to initial gap width per experiment for statistical analysis). (l) Venn-diagram of 4123 with *ELF3* significantly co-expressed genes in the TCGA ccRCC cohort (adjusted p-value <0.0001) and annotation to databases of immune (Immunome), extracellular matrix (Matrisome, including secreted factors) and epithelial–mesenchymal transition (EMTome) related genes. Selected gene list shows secreted factors that are significantly co-expressed with *ELF3*. Colored boxes indicate (co-) annotation of genes to respective databases. Dot plots show independent replicates analyzed (error bars indicate mean and S.E.M.; n.s. – not significant; * p<0.05; ** p<0.01 and *** <0.001).

## Discussion

The large family of annexin proteins show a very heterogeneous cell- and tissue-specific expression pattern.^17^ Here, we identified annexin A4 as specifically enriched in tumor cells of clear cell renal cell carcinoma (ccRCC), in line with previous studies reporting increased protein levels of ANXA4.^23,39^ Given the context- and cell-type dependent role of individual annexins, a comprehensive functional characterization of ANXA4 in ccRCC was to date missing.^17^ In this study, we demonstrated that ANXA4 exhibits distinct subcellular localization patterns related to separate biological function such as membrane repair as well as modulation of transcriptional signatures employing in vitro, in silico and in situ approaches.

ccRCC is the most prevalent subtype of malignant renal tumors and given its dismal prognosis in metastasized patients,^3^ large attempts for better molecular characterization and categorization have been undertaken by integrating various biological layers (e.g. transcriptomic, genomic, proteomic as exemplified by the CPTAC consortium).^8^ Including proteomic, but mostly transcriptomic analysis led also to individual identification of various members of the annexin protein family such as ANXA1 to ANXA5, and more recently ANXA8 and ANXA13 in ccRCC.^18–22^ By utilizing proteomic data, we aimed for a global annotation of annexin proteins which highlighted a specific enrichment of ANXA4 in cRCC (compared to detection levels of other annexin proteins - Figure 1). Integration of scRNA-Seq data moreover validated these findings, while in situ detection confirmed a selective expression pattern of ANXA4 in respective tumor cells. Remarkably, other tumor-enriched annexin proteins such as ANXA2 were not exclusively detected in tumor cells but also in endothelial cells (Figure 1), further supporting the importance of integrative approaches to specify expression patterns. More recently, ANXA8 and ANXA8L were described as enriched proteins in ccRCC.^22^ However, our proteomic annotation did not show significant detection levels, which might be related to detection thresholds or pre-defined cut-off values in respective analytical approaches^25–27^ and does not completely exclude the presence of ANXA8 (as also shown by immunohistochemistry in a previous report).^22^

The majority of investigated annexin proteins in cancer are described to be linked with promoting proliferative behavior.^16^ Remarkably, our studies employing knockdown as well as knockout approaches did not detect significant impact of ANXA4 protein on proliferative behavior of tested cell lines (Figure 2). This is also in contrast to other studies on the role of ANXA4 in other malignancies such as gallbladder carcinoma or ovarian carcinoma, where the authors observed remarkable impact of ANXA4 on proliferation rates.^31,40^ These contrasting observations may relate to employed cell lines (in this study renal carcinoma cell lines such as A498, 786-O or non-malignant proximal tubular epithelial cells, hRPTEC) or may be explained by the general rather low proliferative behavior of ccRCCs (in contrast to respect invasive cancer entities as gallbladder or ovarian carcinoma). However, the molecular mechanism how annexin proteins influence and modulate proliferation (as one key feature of malignant disease) is not yet completely clear and appears to involve rather distinct pathways (e.g. ERK1/2 signal pathways in ANXA3, ANXA5, ANXA6 or b-catenin-dependent pathways in ANXA2).^16^ The underlying mechanism leading to upregulation of various members of the annexin protein family in neoplastic disease is not yet completely understood. Remarkably, our experiments and analysis imply a pVHL-dependent regulation of ANXA4 expression in ccRCC (Figure 2). *VHL*-loss is one of the earliest genetic hallmarks of ccRCC carcinogenesis and is subsequently related to accumulation of HIF1/2 transcription factors.^2^ In this context, previous work showed that also other annexins such as ANXA1 are directly regulated by HIF1alpha or respond to long term hypoxia culture such as ANXA6.^41,42^ Therefore, our observations of pVHL-dependent accumulation of ANXA4 in ccRCC might also indicate a common regulatory response of annexin proteins to hypoxic/HIF-dependent signaling.

Aside from the selective expression of ANXA4 in ccRCC cancer cells (Figure 1), our in situ and in vitro localization studies demonstrated that ANXA4 shows distinct subcellular distribution patterns (Figure 3), which are (at least in vitro) modulated by external stimuli such as acute membrane stress (exerted by hypoosmolarity) or cell densities (Figure S2). Similar observations have been made for ANXA4 for example in colorectal carcinoma, where predominant expression levels of ANXA4 have been linked to a rather aggressive clinical course characterized by increased rates of metastasis.^43^ Interestingly, a nuclear localization pattern was rather exclusively observed in clear cell ovarian carcinoma when compared to other subtypes such as serous or endometrioid ovarian carcinomas.^31^ Our observations in a small cohort of ccRCC cases with either progressive or non-progressive disease course indicated that increased membrane recruitment (and at the same time reduced nuclear localization) of ANXA4 is more prevalent in ccRCC cases with aggressive biology characterized by increased propensities for metastasis (Figure 3). One potential explanation for this predominant membrane recruitment might be related to our observations of ANXA4 dependent membrane repair in renal carcinoma cells (Figure 3). Invasive growth behavior (required for metastasis) is not only depending on degradation of the extracellular matrix, but also challenges respective tumor cells to transmigrate through a dense extracellular environment.^44^ Aside from a dramatic transition in terms of cellular morphology and migration modes (summarized under the concept of epithelial-mesenchymal-transition), also cellular membranes are exposed to high levels of stress.^45^ This has recently led to the concept that acute membrane repair might also be centrally involved in the cascade of essential mechanisms required in metastasis and resistance to cytotoxic immune cells.^46,47^ Our observations that ANXA4 is modulating membrane repair of renal carcinoma cells (Figure 3), is in line with previous studies investigating the general capacity of annexin proteins to reseal and repair damaged plasma membranes (including ANXA6, ANXA5 and ANXA2).^46,48,49^ Remarkably, the potential of ANXA4 for membrane repair appears not to be limited to cancer cells, as it was recently demonstrated that ANXA4 is also involved in the pathogenesis of glaucoma via mediating modes of membrane repair under biophysical strain conditions.^50^ Thereby, these observations complete our understanding of the general involvement of ANXA4 in the process of membrane repair, and further demonstrate the versatile localization modes related to extracellular and tumor inherent features.

With the advent of immune checkpoint-based therapies, the relevance of the tumor immune microenvironment (TIME) has become increasingly apparent.^51^ Utilizing machine-learning based cellular segmentation and cell-proximity analysis in correlation to ANXA4 protein abundance we identified distinct compositional alterations of the TIME in ccRCC. Here, lower levels of ANXA4 protein corresponded to increased immune cell infiltration and more deposited (acellular) extracellular matrix (Figure 4&5). In general, ccRCC is a highly immunogenic tumor characterized by an intricate tumor immune microenvironment (including T-, B-, NK-cells, macrophages and myeloid cells). It is assumed that high levels of tumor infiltrating lymphocytes correspond overall to better outcome, whereas more detailed phenotyping of the TIME recently revealed that polarization of macrophages or angiogenesis signatures more precisely predict TKI-based treatment response on survival.^52^ ECM deposition or peritumoral fibrosis is a rather uncommon feature in ccRCC, whereas recent studies have highlighted that composition of the deposited matrix correlated to survival and progression.^53^ Transcriptome based analysis utilizing our in vitro model of *ANXA4* knockout cells revealed a significant impact on ECM and ECM-related gene sets (Figure 6). While ECM is most likely not solely deposited by tumor cells, but also involves cancer-associated fibroblasts,^54^ our observations imply a potential crosstalk responsible for the observed changes in the TIME depending on ANXA4 levels (Figure 4). Previous studies have implicated NF-κB dependent transcriptional regulation as a potential explanation for transcriptome changes related to ANXA4 function.^31,36,40^ Interestingly, our bioinformatics transcription factor enrichment analysis identified the ETS-family member ELF3 as the most relevant transcription factor for observed transcriptional regulation in our *ANXA4*-knockout model (Figure 6). ELF3 has been described as a major regulator of epithelial differentiation and early mouse knockout studies demonstrated the pivotal importance for intestinal epithelial cell differentiation.^55^ In the context of cancer, more recent studies showed that ELF3 controls processes of epithelial-mesenchymal-transition and thereby impacts tumor features such as migration and invasion.^37,56,57^ In line, with these observations our experiments demonstrate that ELF3 expression recapitulates the impaired migratory mode of *ANXA4* knockout cells (Figure 7). These findings might be explained by a potential EMT-promoting effect of ANXA4, as it was also recently demonstrated for ANXA10 in cholangiocarcinoma.^58^

Together our observations identify ANXA4 as a highly specific marker of cancer cells in ccRCC, exhibiting a versatile subcellular localization pattern which relates to differing cellular functions such as membrane repair as well as co-transcriptional modulation. Our correlative tissue-based segmentation approaches further imply a role of ANXA4 in shaping the tumor composition and microenvironment, which might relate to differing biological behavior. Future studies will be required to dissect the role and complex interplay of individual annexin proteins in ccRCC, and how this knowledge might be further exploited for therapeutic intervention or diagnostic testing.

## Methods

### Cell Culture

Renal cancer cell lines A498 and 786-O (WT and with *VHL* re-expression) as well as human renal proximal tubule epithelial cells (hRPTEC) were used and obtained as recently described.^59^ A498, 786-O and hRPETC cell lines were cultured below 80% confluence using culture medium comprising RPMI-1640 GlutaMAX™ (Thermo Fisher Scientific Inc., Waltham, MA, USA, #61870036) supplemented with 10% FCS (Merck KGaA, Darmstadt, Germany, #S0615) and 1% penicillin-streptomycin (Thermo Fisher Scientific, #15140122). Experiments were done in non-confluent culture conditions, unless stated differently. Cell lines comprising *ANXA4*-knockdown (KD) were transduced with small hairpin RNAs in the pLKO.1 lentiviral vector designed by The RNAi Consortium (TRC). *ANXA4*-KO cell lines were generated using CRISPR/Cas9 technique. Plasmid transduction in order to generate *ANXA4*-KD, *ANXA4*-KO as well as cell lines overexpressing *ANXA4* or *ELF3* and all respective controls was achieved with respective lentiviral particles produced via PEI-transfection in HEK293T/17 cells (ATCC, Manassas, VA, USA, CRL-11268). HEK293T/17 cells were cultured in DMEM GlutaMAX™ (Thermo Fisher Scientific, #31966-021) supplemented with 10% FCS. Routine testing of cell lines for contamination with Mycoplasma was performed using a commercial PCR kit (Mycoplasma PCR detection kit, Hiss Diagnostics GmbH, Freiburg, Germany).

### Expression Plasmids

*ANXA4*-KD: Validated pool of shRNAs targeting *ANXA4* (shRNA-78: GCACACTTCAAGAGACTCTAT, shRNA-80: GCAGAAATTGACATGTTGGAT, shRNA-81: GCCTTGAAGATGACATTCGCT) cloned into pLKO.1 plasmids were purchased (Horizon Discovery Biosciences Limited, Cambridge, UK – TRCN0000056278, TRCN0000056280, TRCN0000056281). Scramble pLKO.1 shRNA control (further termed “scramble”) was a gift from David Sabatini (Addgene plasmid #1864; http://n2t.net/addgene:1864; RRID:Addgene_1864). oeELF3: Human *hELF3* (NM_004433) in pLenti-C-Myc-DDK-P2A-Puro was purchased (OriGene Technologies, Rockville, USA – CAT#: RC200631L3). oeANXA4: Human *ANXA4* (NM_001320698.2) in pcDNA3.1+/C-(K)-DYK vector was purchased from GenScript (GenScript Biotech, Piscataway Township, USA – Clone-ID: OHu26294) and *ANXA4* as well as Luciferase were subcloned into pWPXLd and pLenti-C-Myc-DDK-P2A-Puro expression vectors, respectively; pWPXLd was a gift from Didier Trono (Addgene plasmid #12258; http://n2t.net/addgene:12258; RRID:Addgene_12258).

### CRISPRS/Cas9

To generate *ANXA4* knockout in the A498 kidney carcinoma cell line CRISPR/Cas9 gene editing technology was used. sgRNAs targeting *ANXA4* were designed using the web tool based application CHOPCHOP (version 3 – https://chopchop.cbu.uib.no/). Guide sequences (non targeting – nt-sgRNA-1: 5’-GCGGGCAGAACGACCCTGAC -3’, nt-sgRNA-2: 5’-GAAGACGTGCTGGCGTCACC -3’; ANXA4-targeting – ANXA4-sg1: TCTGCTGGCTATAGGTAAGC(TGG), ANXA4-sg2: GGACGATGCTCTCGTGAGAC(AGG) – PAM sequence in brackets) were cloned into TLCV2-LoxP backbone (TLCV2-LoxP was a gift from Adam Karpf (Addgene plasmid #127098; http://n2t.net/addgene:127098; RRID:Addgene_127098) and respective constructs transformed into competent E.coli (C3040H, New England Biolabs, Ispwich, USA) miniprepped and Sanger sequenced using hU6-F primer (5’-GAGGGCCTATTTCCCATGATT-3’). Target cells were transduced and later selected applying 1 µg/ml Puromycin for 72 h. Further, Cas9 expression from TLCV2-LoxP was induced using 1 µg/ml Doxycycline for 72 h before evaluating knockout efficiency by western blot. Single cell clones of control and *ANXA4* gRNA cell lines were generated and verified by western blot.

### Antibodies and dyes

Following antibodies and dyes were applied for immunofluorescence (IF), Immunohistochemistry (IHC) and western blot (WB) analysis: ANXA4 (HPA007393, Atlas Antibodies, Sweden – WB: 1:1000, IF: 1:300, IHC: 1:300 and sc-46693, Santa Cruz Biotechnology, USA – WB: 1:1000), ANXA2 (#8235, Cell Signaling Technologies, USA – WB: 1:1000, IHC: 1:300), ANXA3 (HPA013398, Atlas Antibodies, Sweden – WB: 1:1000, IHC: 1:300), TUBA (T9026, Merck, WB 1:3000), ELF3 (HPA003479, Atlas Antibodies, Sweden – WB: 1:1000), Luciferase (NB100-1677-100 Novus Biological, Bio-Techne GmbH, Germany), CLDN1 (ab15098, abcam, Cambridge, UK), Hematoxylin, Hoechst 33342, trihydrochloride, trihydrate (H3570, Thermo Fisher Scientific, IF 1:1000), Alexa Fluor phalloidin 488 (A12379, Thermo Fisher Scientific, IF 1:750). For immunofluorescence fluorophore conjugated secondary anti-rabbit antibody (A31572, Thermo Fisher Scientific, IF 1:500) and for western blot HRP-linked, anti-mouse (P0447, Dako, Agilent, CA, USA, WB 1:10000), anti-rabbit (7074, Cell Signaling, WB 1:5000) and anti-goat (P0449, Dako, Agilent, CA, USA, WB 1:5000) antibodies were applied.

### Western Blot Analysis

For western blot analysis, cells were seeded into fresh 10 cm or 6 well cell culture dishes and cultured overnight. Subsequently, cells were scraped in PBS and pelleted by 100 g centrifugation for 5 min at room temperature (RT) and supernatant was discarded. The cell pellet was resuspended and lysed in RIPA buffer for 15 min followed by 15 min centrifugation at 16000 g at 4°C. The supernatant was collected and mixed with a 2× Laemmli sample buffer containing DTT and denatured at 95°C for 10 min. Protein concentration was determined using Pierce BCA Protein assay kit (#23225, Thermo Fisher Scientific) and loading equalized accordingly. Cell lysates were analyzed by SDS-polyacrylamide gel electrophoresis (SDS-PAGE) and western blotting applying the trans-blot turbo transfer system (Bio-Rad Laboratories, Inc., Hercules, CA, USA) and PVDF membranes (#1704157, Bio-Rad). Membranes were blocked in 5% BSA in TBS-T and primary antibodies incubated overnight at 4°C. Following 3× washing steps of 5 min in TBS-T membranes were incubated for 1 h at RT in respective HRP-linked secondary antibody. After 3× washing of 5 min in TBS-T antibody detection was performed by HRP-ECL chemiluminescence reaction (#32109, Thermo Fisher Scientific) using the Fusion FX 7 (Vilber Lourmat GmbH, Germany) chemiluminescence imager. Densitometry analysis was performed in Fiji ImageJ v1.54f as the ratio of the target signal to the respective TUBA (alpha-Tubulin) band. Values were normalized to the mean of respective control measurements per sample and experiment for graphical depiction and statistical analysis.

### Transwell assays

For assessment of cellular migratory and invasive capacity Transwell/Boyden-chamber-assays were used with respective pore diameter and uncoated or matrigel-coated membrane. Transwells were placed in 24-well plates (662160, Greiner Bio-One, Kremsmünster, Austria). In order to analyze migration 8 µm pore membrane transwells were used (662638, Greiner Bio-One, Kremsmünster, Austria). To analyze invasion the same 8 µm pored transwells were coated with in PBS diluted growth-factor reduced matrigel (356231, Corning, New York, USA) in a final concentration of 0.3 mg/ml and incubated for 1h at 37°C. Remaining non-polymerized diluted matrigel was discarded before cell seeding. For all transwell assays cells were trypsinized for 5-10 min, centrifuged for 5 min at 100 g and washed once in PBS before resuspending in serum-free culture medium followed by cell counting using a hemocytometer. Cells were diluted to 250.000 cell/ml and seeded into the upper chamber in 400 µl. Afterwards the lower chamber was filled with 800 µl of regular culture medium (supplemented with 10% FCS) followed by incubation in 37°C. Incubation time was dependent on the assay and cell line used (A498: migration 6 h; invasion: 18 h). After incubation, cells on the transwell membrane were fixated in 4% PFA in (Electron Microscopy Sciences, #15714-S) diluted in PBS (Thermo Fischer, #10010023) for 20 min, added to top and bottom chamber. After fixation, transwells were washed 3× in PBS and cell nuclei stained using Hoechst for 30 min in darkness. Cells which did not migrate through the transmembrane were removed by scraping gently with cotton swabs (XL56.1, Carl Roth GmbH, Karlsruhe, Germany) on top of the transwell membrane. After subsequent washing 3× in PBS cells which migrated on the bottom of the transwell were directly imaged using an Axio Observer (Carl Zeiss AG, Oberkochen, Germany) 10× magnification. Images were stitched in Zen blue 3.0 software (Carl Zeiss AG, Oberkochen, Germany) and cells on the bottom of the transwell counted using QuPath version 0.5.1.^60^ Counted cells were normalized to the respective area analyzed and graphically depicted as the number of cells per cm².

### Gap closure / collective cell migration

To assess the migratory capability of cells in a collective cell cluster gap closure assays using culture-insert dishes (81176, Ibidi GmbH, Gräfelfing, Germany) were performed. 50000 cells are seeded in 70 µl of regular culture medium into each well of the insert, respectively, and incubated overnight at 37°C incubator to enable cell adhesion and formation of a confluent cell layer. By pulling of the insert, using a sterile forceps an approximate gap of 500 µm was formed. The cells were subsequently washed once with PBS and 2 ml of regular culture medium was added. Phase contrast images of the gap are taken with an Axio Observer microscope (Carl Zeiss AG, Oberkochen, Germany) using 5× magnification at respective time points commencing with 0 h directly after gap creation. Gap width at every time point was further measured using the wound size healing tool in Fiji ImageJ v1.54f.^61^ Graphs are depicted as the relative gap width to the respective gap width at the starting point 0 h after gap creation.

### Immunohistochemistry

To stain target proteins *in situ* immunohistochemistry (IHC) was used. Patient tumor samples were fixated in 4% formaldehyde solution. After embedding the tissue in paraffin, 2 µm slices were cut using a microtome, transferred to microscope slides and dried overnight at 65°C. After deparaffinization samples did undergo heat-induced antigen retrieval (HIAR) for 5 min in pH 6 citrate buffer using a pressure cooker. Afterwards samples were washed with PBS, endogenous peroxidases inactivated by application of 3% H_2_O_2_ solution for 10 min and after washing once more with PBS, blocked using 5% BSA in PBS for 1 h at RT. Subsequently, respective primary antibody was diluted accordingly in 5% BSA in PBS and applied for 1 h at RT. After further washing 3× with PBS anti-rabbit antibody EnVision FLEX+ (K8009, Dako, Agilent, CA, USA) was applied for 30 min. The final washing steps of five times with PBS were followed by application of Liquid DAB+ Substrate Chromogen System (K3468, Dako, Agilent, CA, USA) to visualize bound antibodies. Slides were counterstained using Hematoxylin and sealed by coverslips. Stained slides were digitalized using a Ventana DP 200 slide scanner (Roche Diagnostics Deutschland GmbH, Mannheim, Germany) and analyzed in QuPath (version 0.5.1).

### Nuclear localization scoring

Samples from primary ccRCC tumors of a cohort of matched patients with progressive (metastasized) or non-progressive disease have been gathered and processed in accordance with the IHC protocol outlined in the preceding section (see Supplementary Spreadsheet S4 for cohort characteristics). Subcellular localization of ANXA4 was classified in 4-tier scores for respective subcellular compartment (nucleus, cytoplasm, plasma membrane). Nucleus localization score: 0 – nuclear exclusion: ANXA4 is not detectable within the cell nucleus; 1 – weak nuclear localization: ANXA4 can be distinguished within the nucleus to a small extent; 2 – medium nuclear localization: ANXA4 is detectable in the nucleus; 3 – strong nuclear localization: ANXA4 localizes primarily within the nucleus depicted by strong nuclear staining. Cytoplasm localization score: 0 – ANXA4 not detectable within cytoplasm; 1 – weak staining of ANXA4 in cytoplasm; 2 – intermediate ANXA4 staining in cytoplasm; 3 – strong ANXA4 staining in cytoplasm. Plasma membrane localization score: 0 – ANXA4 does not localize to the plasma membrane: plasma membrane is indistinguishable 1 – ANXA4 does localize to the plasma membrane: plasma membrane can be distinguished; 2 – ANXA4 distinctly localizes to the plasma membrane; 3 – ANXA4 dominantly localizes to the plasma membrane.

### Immunofluorescence, cellular stressor application

For immunofluorescence (IF) analysis cells were seeded on ibiTreat-8well-dishes (80826, Ibidi GmbH, Gräfelfing, Germany) for IF under stress or density conditions. Cellular stress conditions were applied as follows. For digitonin, a pore forming detergent, digitonin (D141, Sigma-Aldrich, St. Louis, USA) was diluted in regular culture medium to a final concentration of 20 µM and applied for 10 min. For hypoosmolaric conditions regular culture medium was diluted with ddH_2_O to reach an osmolarity of 25 mOsm/l (RPMI 1640 without supplements: 260 – 310 mOsm/kg – according to manufacturer) and applied onto the cells for 15 min. For all experimental approaches cells were washed with PBS once and fixated in 4% PFA (Electron Microscopy Sciences, #15714-S) in PBS (Thermo Fischer, #10010023) for 20 min. Cells were further washed 3× with PBS and permeabilized using 0.1% Triton X-100 diluted in PBS for 5 min. After washing 3× with PBS cells were blocked with 5% BSA in PBS for 1 h at RT, proceeding with incubation of respective primary antibody in 5% BSA in PBS overnight. Cells were then washed 5× with PBS and incubated with respective secondary antibody for 1 h at RT before fluorescence imaging using an Axio Observer microscope (Carl Zeiss AG, Oberkochen, Germany) equipped as described before.^62^

### Plasma membrane permeability assay

To assess the integrity of the plasma membrane under membrane stress conditions 3000 cells/well were seeded in regular culture medium in 8well-dishes (80826, Ibidi GmbH, Gräfelfing, Germany) and settled over night. Cells were placed in a controlled climate chamber setup (37°C, 5% CO_2_) of Zeiss AxioObserver microscope (Carl Zeiss AG, Oberkochen, Germany) and allowed to acclimatize for 15 min while image regions were selected and focus points adjusted. Medium was removed and culture medium supplemented with 0.2 µg/ml impermeable Hoechst 33258, Pentahydrat (bis-Benzimid) (H3569, Thermo Fisher Scientific Inc., Waltham, MA, USA) and 20 µg/ml digitonin (D141, Sigma-Aldrich, St. Louis, USA), a pore-forming detergent, acting as membrane stressor. Dye uptake into cells with damaged plasma membrane was measured over time using mentioned Zeiss AxioObserver microscope. Before addition of stressor/dye-solution and after the last time point phase-contrast pictures were taken to obtain total cell numbers. Pictures were analyzed by segmentation, thresholding and counting of cells with stained nuclei compared to total number of cells in image regions using Fiji ImageJ v1.54f.

### BrdU assay

To assess cell proliferation BrdU ELISA was performed (BrdU Cell Proliferation ELISA Kit (colorimetric), ab126556, abcam, Cambridge, UK) following protocol instructions. Cells were seeded at 2 x 10^5^ cells/ml in 200 µl/well of regular culture medium with respective controls (blank: medium only, background: cells without BrdU) in 96-well plate. 20 µl of diluted 1× BrdU were added to appropriate wells and incubated for 18 h. Cells were fixed and DNA denatured using Fixing Solution and washed with before addition of anti-BrdU monoclonal Detector Antibody for 1 h. After washing as before, 1× Peroxidase Goat Anti-Mouse IgG Conjugate was freshly prepared and added for 30 min. Following a final wash with Wash Buffer the plate was flooded with ddH_2_O, dried on paper towels and TMB Peroxidase Substrate was added for 30 min. The reaction was stopped using Stop Solution and the plate measured at dual wavelength of 450 nm / 550 nm. 450 nm intensity is proportional to the amount of BrdU incorporated in proliferating cells, while 550 nm is out of range of the reaction product resembling noise and is subtracted from the 450 nm value.

### Incucyte growth assay

To assess cellular proliferation the Incucyte S3 Live-Cell Analysis System (Sartorius, Göttingen, Germany) was used to obtain growth curves with a high sensitivity. Cells were seeded in 96-well plates, 1000 cells in 200 µl per well of regular culture medium and observed for 7 days taking pictures every 2 h. Analysis of cellular growth was conducted using the built-in Incucyte software calculating the cell confluence over time in the pictures taken. Graphs depict the relative confluence of cells in the pictures taken.

### Transcriptome analysis

RNA Sequencing analysis was performed as recently described.^63^ In brief, equalized numbers of *ANXA4* cell lines (Ctrl.-1, Ctrl.-2, KO-1, KO-2) were seeded into cell culture dishes and cultured applying standard cell culture conditions for 24 hours. Cells were harvested by scraping and total RNA was isolated using the Monarch Total RNA Miniprep kit (T2010S, New England Biolabs Inc., Ipswich, MA, USA) according to the supplierś instructions. RNA of two independent replicates was isolated. RNA quality was measured as RQN >9.3 for all samples using an Agilent Fragment Analyzer. Following processing steps (Poly(A) mRNA selection, library preparation (NEBNext Ultra II RNA Library Prep kit, New England Biolabs Inc.) and Illumina NovaSeq 2 × 150 bp paired-end sequencing) were performed by Azenta Life Sciences, Leipzig, Germany. A minimum of 19.8 million reads per samples were measured. The Galaxy Europe resource was used for further processing and analysis of sequencing data.^64^ In brief, reads were processed by fastp, HISAT2 was applied for read alignment, featureCounts and DESeq2 were used for differential gene expression analysis and DESeq2 was used for principle component analysis (PCA). The GenePattern platform was used for gene set enrichment analysis (GSEA) for GO-Terms.^65^ ChEA3 was used for transcription factor enrichment analysis (TFEA).^66^ The Matrisome DB, Innate DB and EMTome DB resources were used for gene annotation.^67–69^ See Supplementary Spreadsheet S6 for transcriptome and gene enrichment analysis. Sequencing data have been deposited in the NCBI Gene Expression Omnibus and are accessible via GEO series accession numbers GSE267754.

### Proteomics analysis

A498 ctrl., A498 +VHL, 786-O ctrl. and 786-O +VHL cells were seeded and cultured at standard conditions for 48 hours. Proteome analysis of cell lines was performed as previously described.^70^ See Supplementary Spreadsheet S3 for analysis of proteome data. All datasets are available at the MassIVE repository (http://massive.ucsd.edu/; dataset identifier: MSV000094814).

### Analysis of ccRCC cohorts and resources

Transcriptome and proteome expression data of the annexin protein family in ccRCC tumor tissue versus normal adjacent tissue (NAT) were retrieved form datasets of three previously published ccRCC cohorts (see Supplementary Spreadsheet S1 for details).^25–27^ Single cell RNA sequencing (scRNA-Seq) data were assessed and analyzed via the CELL×GENE platform from a previously published cohort of 7 ccRCCs.^28,29^ Expression of *ANXA4* in single cell types was mapped to predefined t-SNE clusters and clipped to percentile ranges via the CELL×GENE explorer. Analysis of annexin family and marker genes, relative gene expression scaling, annotation to predefined cell types and dot blot generation was performed via the CELL×GENE gene expression tool.

The cBioPortal for Cancer Genomics was used for analysis of the TCGA Kidney Renal Clear Cell Carcinoma (TCGA, Firehose Legacy) and Cancer Cell Line Encyclopedia (Broad, 2019) datasets.^7,30,71^ For survival analysis, best gene expression (FPKM) cut-offs for survival between patient groups in the TCGA cohort were previously calculated and used to define *ANXA4* high and *ANXA4* low ccRCCs accordingly.^72^ Kaplan-Meier survival analysis and caparison of clinical parameters between these two patient groups as well as calculation of statistical characteristics was performed using the cBioPortal for Cancer Genomics toolbox (Supplementary Spreadsheet S2). In addition, mRNA co-expression analysis for *EFL3* in the Kidney Renal Clear Cell Carcinoma (TCGA, Firehose Legacy) cohort was retrieved from the cBioPortal for Cancer Genomics platform. The Matrisome DB, Innate DB and EMTome DB resources were used for annotation of *ELF3* co-expressed genes (see Supplementary Spreadsheet S7 for co-expression analysis of *ELF3*).^67–69^

For analysis of renal cancer cell lines, the Cancer Cell Line Encyclopedia (Broad, 2019) was filtered for renal cancer cell lines and gene expression z-scores, *VHL* mutation profiles and segmented copy number (CN) data for chromosomal segments were retrieved. Loss of the chromosome 3p arm was estimated by underrepresentation of respective CN segments (mean log_2_ FC <-0.5 for 3p segments). See Supplementary Spreadsheet S1 for details.

Transcriptome and proteome data, cancer characteristics and ESTIMATE scores of the ccRCC CPTAC cohort were retrieved from the published datasets.^27^ Z-scores of log_2_ transformed FPKM transcriptome and of ln transformed DIA proteome data from 213 cancer tissue samples were calculated and used for selection of 26 ANXA4 high and 18 low ccRCC patients as well as analysis of gene expression profiles in these samples (see Supplementary Spreadsheet S5 for details). Images of corresponding HE stained FFPE tissue samples were download from the CPTAC-CCRCC dataset from the Cancer Imaging Archive. Cases that only contained images of cryopreserved samples were excluded from further image analysis. Subsequent image analysis of the remaining 24 ANXA high and 18 ANXA low cancers was performed using the QuPath v0.4.3 software.^60^ In brief, whole tumor tissue sections were manually annotated as analysis region. Areas that were out of the focus plane were excluded from further analysis. Cancer tissue composition was automatically determined by sample adapted supervised training of a pixel classifier using the random trees machine-learning algorithm implemented into QuPath (1.98 µm/pixel resolution, using all deconvolutions of image channels and Gaussian-, Laplacian-, weighted deviation-, gradient magnitude- and structure tensor maximum eigenvalue pixel features for classification). Thereby, pixel and sample regions were classified as tumor tissue, stroma tissue, erythrocyte filled space, serum filled space or empty space. The StarDist U-Net was applied for cell segmentation.^73^ After quality control, shape and staining features were calculated for all segmented objects (cells). These features were used for sample adapted supervised training of an object classifier using the machine-learning algorithm implemented in QuPath. Cells were classified as Lymphocyte (LC), tumor cell (TC), stroma cell (SC) or other cell (e.g. granulocytes and image artefacts). Other cells were eliminated from subsequent analyses and calculation of cell-type proportions. Lymphocyte hotspots/clusters were annotated using the density map tool (50 µm radius). Cell network features were calculated applying delaunay cluster analysis (15 µm threshold) and the object centroid distance plugin for cell classes. The CytoMAP 1.4.21 toolbox was used for analysis of cellular regions.^74^ Therefore, objects (detected cells) including X-Y coordinates and classifications were loaded into CytoMAP. Cell neighborhoods were calculated (50 µm radius) and neighborhoods were clustered into 8 regions by an self-organizing map algorithm using cellular composition and density as cluster parameters. These cell cluster regions and cluster composition were used for subsequent analysis. See Supplementary Spreadsheet S5 for results of histological analysis of CPTAC samples.

### Statistical analysis

For statistical analysis and graphical depiction of experimental data, the GraphPad Prism 8 software was used. Scatter plots indicate respective individual units (replicates, samples) used for statistical testing while bars or single dots indicate mean values of replicates or samples, described accordingly in detail within respective figure legend. Error bars indicate the standard error of the mean (S.E.M.) or confidence interval (CI) as indicated. Statistical tests were conducted based on the respective experimental design and data distribution. Statistical analysis of Figure 1h, 1i, 1j, 1k was performed using the cBioPortal platform as was described in the respective methods section. Unpaired student’s t-test (Figure 2d, 2e, 2j, 3h, 4k, 5a, 5e, 7d, 7f, 7i, 7k), ordinary one-way ANOVA combined with Tukey’s multiple comparisons test (Figure 2k, 3i, 4g, 4n, 7b, 7c), Two-way ANOVA with Sidak’s multiple comparisons test (Figure 3j), unpaired student’s t-test followed by calculation of the false discovery rate (FDR) using the method of Benjamini, Krieger and Yekutieli (Figure 4d), unpaired t test with Welch’s correction (Figure 4h, 4i, 4j, 4l, 5b, 5c, 5i, 5j) and Pearsons correlation analysis (Figure 4m, 5d) were used for presented analyses. Statistical analysis of transcriptome and proteome data as well as implementation of analyses from external databases is described in the respective methods sections. Statistical significance was defined as p < 0.05 and significance levels are indicated as * p < 0.05, ** p < 0.01, *** p < 0.001, **** p < 0.0001 and non-significant (n. s.) in respective figure panels. The number of independent experiments and analyzed units are stated in the figure legends and/or respective methods section.

## Acknowledgments

We thank Katja Gräwe, Alena Sammarco and Marlene Schmid for expert technical assistance. In addition, we would like to express our gratitude to all members of our laboratories for helpful discussions and support. The authors acknowledge the support of the Freiburg Galaxy Team: Björn Grüning, Bioinformatics, University of Freiburg (Germany) funded by the Collaborative Research Centre 992 Medical Epigenetics (DFG grant SFB 992/1 2012) and the German Federal Ministry of Education and Research BMBF grant 031 A538A de.NBI-RBC. We thank the Lighthouse Core Facility staff of the Medical Center - University of Freiburg for help with their resources and their excellent support. The Lighthouse Core Facility is funded by the Medical Faculty, University of Freiburg (Project Numbers 2021/A2-Fol; 2021/B3-Fol).

## Author Contributions

Conceptualization, M.R. and C.S.; formal analysis, M.W., M.R., O.S., C.S.; investigation, M.W., M.R., C.G.P., A.M.,T.F., A.L.K., O.S., C.S.; methodology, M.W., M.R., T.F., M.G:, O.S., C.S.; resources, I.F., M.W; project administration, O.S. and C.S.; supervision, M.R., M. Werner, M.G., O.S. and C.S.; validation, M.W., M.R. and C.S.; visualization, M.R. and C.S.; writing—original draft, M.R., M.W. and C.S.; writing—review and editing, M.R. and C.S. with input from all authors; funding acquisition, O.S. and C.S.. All authors have read and agreed to the published version of the manuscript.

## Institutional Review Board Statement

The study was conducted according to the guidelines of the Declaration of Helsinki and approved by the Institutional Ethics Committee of the University Medical Center Freiburg (EK 21-1288). Informed consent was obtained from all subjects involved in the study.

## Data Availability Statement

The data presented in this study are available in the article or supplementary materials. Sequencing data have been deposited in NCBI Gene Expression Omnibus and are accessible via GEO series, accession numbers GSE267754. The Proteomic data is available at the MassIVE repository (http://massive.ucsd.edu/; dataset identifier: MSV000094814).

## Supplementary material

Supplementary Figure S1: Expression analysis of annexin genes corresponding to main Figure 1&2.

Supplementary Figure S2: Cell density dependent localization analysis of ANXA4.

Supplementary Spreadsheet S1: Expression analysis of annexin genes in ccRCC tissue and renal cancer cell lines.

Supplementary Spreadsheet S2: Survival analysis for *ANXA4* expression using the TCGA ccRCC dataset.

Supplementary Spreadsheet S3: Analysis of the pVHL dependent proteome in 786-O and A498 cell lines.

Supplementary Spreadsheet S4: Characteristics of the ccRCC cohort used for IHC analysis of ANXA4.

Supplementary Spreadsheet S5: Histological analysis of ANXA4 high and low ccRCC in the ccRCC-CPTAC cohort.

Supplementary Spreadsheet S6: RNA sequencing analysis of *ANXA4* control (WT) and KO A498 cells.

Supplementary Spreadsheet S7: Co-expression analysis for *ELF3* in the ccRCC-TCGA dataset.

