## Supplementary Figures for "Versatile roles of Annexin A4 in ccRCC: impact on membrane repair, transcriptional signatures, and composition of the tumor microenvironment"

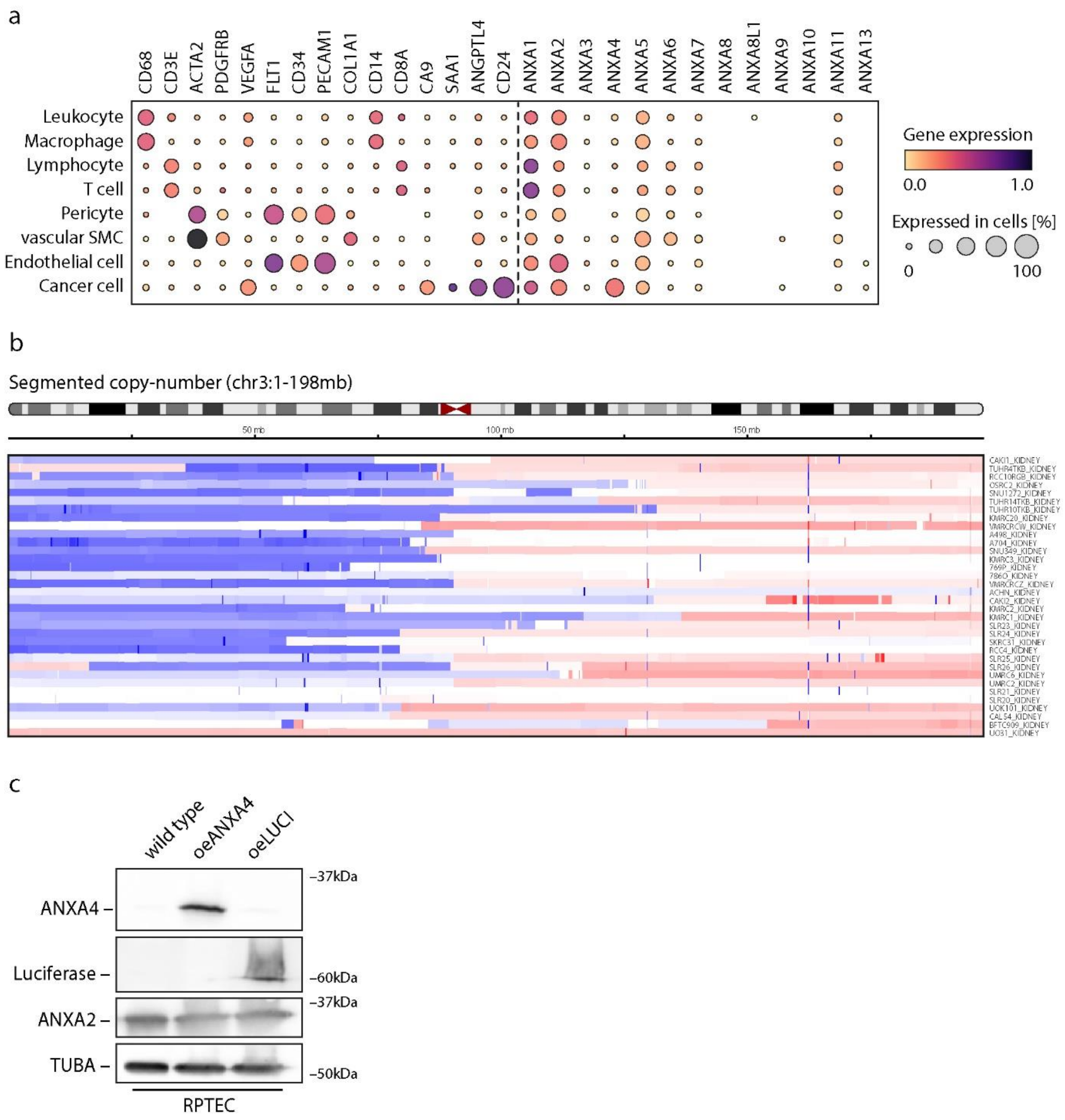


### Supplementary Figure 1

(a) Dot plot indicates relative expression of marker proteins for cell types and annexin family genes in indicated cell populations derived from previously published scRNA-Seq analysis of seven ccRCCs.[1] Color scaling of relative gene expression differ compared to main Figure 1 due to addition of highly expressed cell type markers. (b) Graph shows relative copy-number (CN) changes of segmented chromosome 3 for analyzed renal cell cancer cell lines (mb – mega base; color scale indicates loss (blue) to no alteration (white) to enrichment (red) of $\log_2$ CNs relative to chromosome 3 segments). Analysis and graph were generated via the cBioPortal for Cancer Genomics (Cancer Cell Line Encyclopedia dataset).[2,3] CN changes suggesting loss of CN segments of the p-arm of chromosome 3 (chr3p) in most renal cancer cell lines. (b) Western blot analysis of RPTEC cell lines overexpressing ANXA4 (oeANXA4) or Luciferase (oeLUCI).

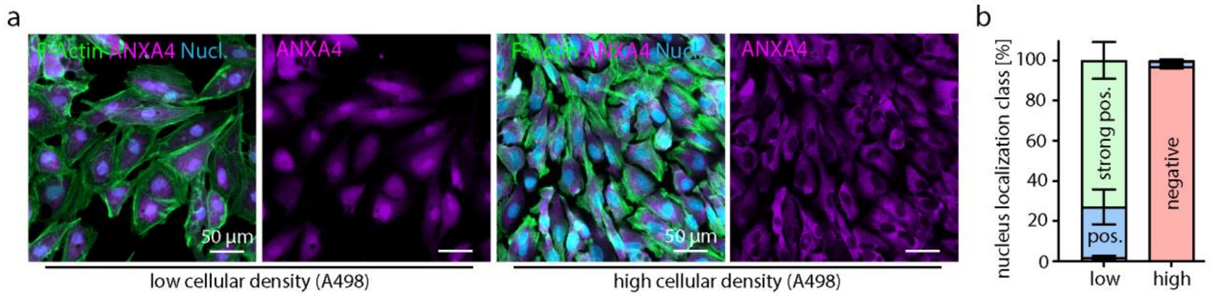


**Supplementary Figure 2**

(a&b) Cell density influences nuclear localization of ANXA4. Representative immunofluorescence images show subcellular localization of ANXA4 at low and high cell density (cells were co-stained by Phalloidin (F-Actin) and Hoechst (Nucleus)). Stacked bar charts show quantification of main subcellular ANXA4 localization into three classes – nucleus negative (red), nucleus positive (blue), nucleus strong positive (green).
